# Systemic signalling through *TCTP1* controls lateral root formation in Arabidopsis

**DOI:** 10.1101/523092

**Authors:** Rémi Branco, Josette Masle

## Abstract

As in animals, the plant body plan and primary organs are established during embryogenesis. However, plants have the ability to generate new organs and functional units throughout their whole life. These are produced through the specification, initiation and differentiation of secondary meristems, governed by the intrinsic genetic program and cues from the environment. They give plants an extraordinary developmental plasticity to modulate their size and architecture according to environmental constraints and opportunities. How this plasticity is regulated at the whole organism level is still largely elusive. In particular the mechanisms regulating the iterative formation of lateral roots along the primary root remain little known. A pivotal role of auxin is well established and recently the role of local mechanical signals and oscillations in transcriptional activity has emerged. Here we provide evidence for a role of Translationally Controlled Tumor Protein (TCTP), a vital ubiquitous protein in eukaryotes. We show that Arabidopsis AtTCTP1 controls root system architecture through a dual function: as a general constitutive growth promoter locally, and as a systemic signalling agent via mobility from the shoot. Our data indicate that this signalling function is specifically targeted to the pericycle and modulates the frequency of lateral root initiation and emergence sites along the primary root, and the compromise between branching and elongating, independent of shoot size. Plant TCTP genes show high similarity among species. TCTP messengers and proteins have been detected in the vasculature of diverse species. This suggests that the mobility and extracellular signalling function of *AtTCTP1* to control root organogenesis might be widely conserved within the plant kingdom, and highly relevant to a better understanding of post-embryonic formation of lateral organs in plants, and the elusive coordination of shoot and root morphogenesis.

## Introduction

Plant development is highly plastic. This is essential to survival and adaptation to a wide range of environments from which, being sessile, plants cannot escape. That plasticity manifests itself as an extraordinary capacity of a plant to modify the number, size, shape, patterning and spatial deployment of its organs, above and below ground, to efficiently adapt to environmental constraints.

As is typical of dicotyledonous species, the Arabidopsis root system arises from a primary root, initiated in the embryo, and *de novo* organogenesis of secondary and higher order lateral roots, post-embryonically [1, 2]. Lateral roots constitute the major part of the root system, and are major determinants of its ability to take-up water and nutrients and to further expand into new soil pockets. Despite their high agronomic and ecological relevance, the molecular mechanisms that determine the placement of LRs, in space and time, and their number are still little known. LR roots originate from inner root pericycle founder cells, through a pre-patterning (priming) activation and cell fate redefinition process that gives them competence to divide and differentiate in an orderly fashion to generate a highly organised root primordium [1, 3–5]. The establishment of LR initiation sites and subsequent actual initiation process are not well-understood. They are thought to involve an oscillating transcriptional network in interaction with auxin and other as yet unidentified mobile signals and, consistently to critically depend on intercellular connectivity [4–6].

There is intense trafficking of a vast array of molecules between shoot and roots – photo assimilates and a range of growth enabling metabolites, numerous hormones, and also a vast number of proteins and RNAs [7–11]. Besides specialised microRNAs (miRNAs) and small interference RNAs (siRNAs), translocated RNAs include a huge cohort of protein encoding mRNAs of various kinds. It is now well-recognised that hundreds to thousands mRNAs transit in the phloem and have long distance mobility between aerial organs and roots [11–15]. Long distance movement of proteins has also been demonstrated, many of which can be unloaded in the root tip [16, 17]. An emerging consensus is that this movement does not simply reflect passive diffusion and mass flow, but in part an actively controlled movement, likely serving the delivery of systemic signalling agents [17–22]. There are a few established cases of genes regulating plant development through mobility of their mRNA and /or encoded protein, such as FT in the regulation of flowering time [23, 24], GIBBERELLIC ACID-INSENSITIVE (GAI) or Mouse eras (me) in the regulation of leaf development [25, 26], *StBEL5* transcription factor in the control of tuber formation [27], or shoot-derived *Auxin/Indole-3-Acetic Acid IAA18* and *IAA28* in the regulation of lateral root formation [28]. However, nothing is known of the physiological significance of the vast majority of mobile mRNAs or proteins transiting through the phloem and roles in receiving cells, nor of the mechanisms controlling their excretion, transport and delivery.

Among the large number of mRNAs with demonstrated long distance mobility between above- and below-ground organs are transcripts encoding Transcriptionally Controlled Tumour Proteins (TCTP). These include Arabidopsis *AtTCTP1* [11, 12] and *AtTCTP2* [29], a *Vitis vinifera* TCTP (*GSVIVG01017723001* [13]), or *Csa3M154390*, a *Cucumis* TCTP [30]. TCTP is a highly conserved ubiquitous protein found in almost all eukaryotes. Its molecular function is still a matter of debate, but clearly relates to the regulation of GTPase activity [31, 32]. Fitting with this, TCTP is involved in a number of fundamental biological processes. It is known as an essential mitotic factor and a promoter of cellular growth interacting with the protein synthesis machinery, and also as a cytoprotective and anti-apopoptic protein (reviewed by [33, 34]). Although best characterised in animals given their high relevance to malignancy and cancer progression, these core functions appear largely conserved in other eukaryotes, including plants, and make TCTPs essential proteins to embryogenesis and early development, organ patterning, regulation of organ size and cellular homeostasis [34]. In Arabidopsis, knock-out mutations of either *AtTCP1* or its homologue, *AtTCTP2* are lethal [35, 36], as is the case of TCTP loss of function in mice [37] or drosophila [31]. Reduced *AtTCTP1* expression through RNA interference causes general cell proliferation and growth inhibition, in both vegetative and reproductive organs [35, 36] and plant TCTPs have been linked to resistance to various abiotic stresses, including salinity, drought, flooding, sub-optimal temperatures [38–43], and also to biotic stresses [44–46].

In addition to its core functions at the cellular level, mammalian TCTP has long been known to act as an extracellular protein in the immune system and was in fact first characterised as a histamine-releasing factor (HRF) [47]. Human TCTP has since been shown to modulate the release of cytokines and other signalling molecules [48] and have a broad role in immunity (reviewed in [49]). Whether plant TCTPs also assume non-cell autonomous functions is unknown. The demonstrated long distance translocation of TCTP transcripts and proteins between scion and root-stock in various species would support that possibility, but in itself does not prove it. Another most interesting indication in that direction is the earlier finding by Aoki and colleagues [50] that pumpkin TCTP (CmaCh11G012000) moved rootward in a destination-, selectively controlled manner when introduced in rice sieve tubes, and furthermore in complex with RNA binding proteins and the conserved eukaryotic translation initiation factor eIF5A. Moreover, this association was found to be necessary to the selective movement of the protein complex.

Together these observations raise the prospect that plant TCTPs might have physiologically important systemic signalling functions, through mobility. This is what we sought to examine, focusing on Arabidopsis *AtTCTP1* [35]. We asked whether long-distance movement of *AtTCTP1* gene products occurs under physiological conditions and plays a role in shaping root architecture.

## Results

### *AtTCTP1* mRNA moves through shoot-root graft junction, in both directions

To investigate long distance mobility of endogenous *AtTCTP1* mRNA and encoded protein in a physiologically relevant context, and be able to differentiate between locally expressed *AtTCTP1* and *AtTCTP1* originating from distant sources, we performed reciprocal grafts between WT (Col-0) and a TCTP1-GFP line expressing a AtTCTP1-GFP fusion protein under the control of *AtTCTP1* native promoter (*pAtTCTP1∷gAtTCTP1-GFP* [35], Fig. 1A). To avoid potential confounding effects from the uncontrolled formation of adventitious roots post-grafting, we developed a modified micro-grafting technique where none is formed (see Methods). Fourteen days after grafting (14 DAG), the scions had developed 5 to 6 leaves of normal size (Fig. 1B), indicating that the scion-root stock junction was fully functional. Microscopic observation confirmed a clean junction, with continuous vasculature and absence of adventitious root primordia (Fig. 1C-F).

**Fig. 1.**
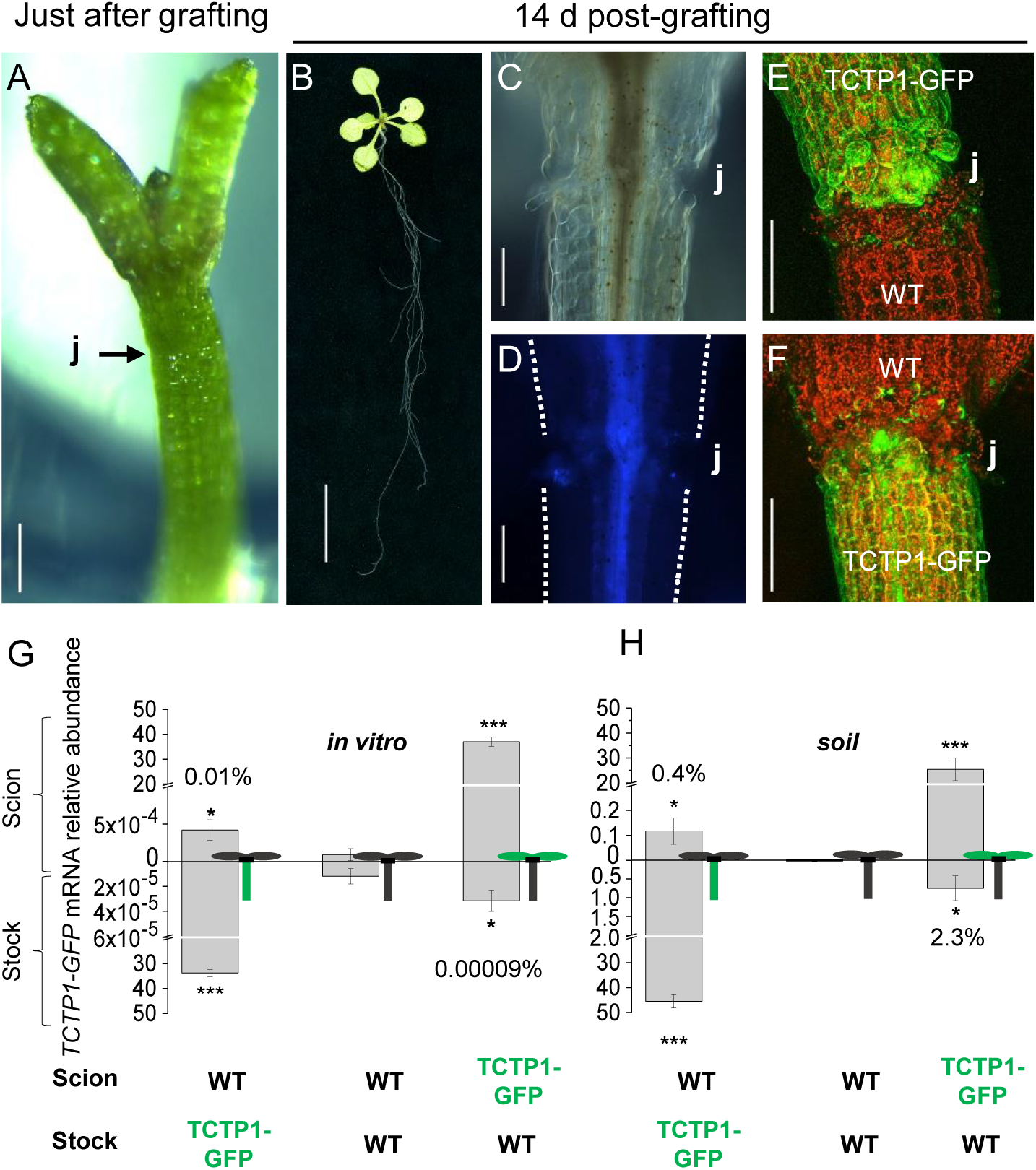
*AtTCTP1* mRNA moves over hypocotylar graft junctions in young and adult plants, in variable proportions. **A-D,** Representative images of: scion-root stock junction (j) immediately after grafting (**A**); whole seedling (**B**) and graft junction (**C-D**) 14 DAG. DIC image (**C**) and autofluorescence image (**D**). Scale bars, 250 µm **(A),** 10 mm (**B**), 200 µm (**C** and **D**). **E** and **F**, TCTP1-GFP fluorescence at the graft junction imaged by confocal microscopy. WT root grafted to *pTCTP1∷gTCTP1-GFP* scion (**E**) and WT scion grafted to *pTCTP1∷gTCTP1-GFP* root (**F**). Scale bar, 200 µm. **G** and **H**, *AtTCTP1-GFP* and endogenous *AtTCTP1* transcripts were quantified by qRT-PCR, in the scion (values above x-axis) and root-stock (values below x-axis) of reciprocal grafts between WT and transgenic *pTCTP1∷gTCTP1-GFP* seedlings grown *in vitro* and sampled at 14 DAG (**G**) or transferred to soil 10 DAG and sampled at 80 DAG (**H**). Scion-root stock combinations are indicated below each panel, and schematised beside each bar. Values are means ± S.E. of fold-change in target gene expression relative to expression of four reference genes (*in vitro* seedlings: n= 5 biological replicates each consisting of pooled rosettes or root-stocks from 5 plants; soil-grown plants: n ≥ 6 biological replicates, each consisting of pooled rosettes or root-stocks from 2 plants). Asterisks denote statistically significant expression differences compared to WT / WT control homograft by one tail Student’s T-test (* *P*< 0.05; *** *P*< 0.001). Similar results were obtained with two sets of GFP-specific primers as listed in Methods.

*TCTP1-GFP* transcripts were detected in both the scion of WT / TCTP1-GFP grafts and the rootstock of TCTP1-GFP / WT reciprocal grafts. This was observed in young seedlings grown *in vitro* (14 DAG, Fig. 1G) and also at a much later stage (early flowering) in soil-grown grafts (80 DAG, Fig. 1H). Mobile *TCTP1-GFP* transcripts were more abundant in the latter, both in absolute terms and relative to endogenous *AtTCTP1* transcripts in the same tissue (0.4% and 2-3% in scion and rootstock, respectively in soil-grown plants compared to 0.001% and 0.00009% in seedlings on agar media). These data demonstrate sustained bi-directional mobility of *AtTCTP1* mRNA, of variable magnitude.

### AtTCTP1 protein of scion origin is detected in WT rootstock, with preferential accumulation in phloem-pole pericycle cells at sites of lateral root formation

We next examined the presence of the encoded TCTP1-GFP protein in these grafts by confocal laser microscopy. A weak TCTP1-GFP fluorescence signal was consistently detected in the primary root of *pAtTCTP1∷g*TCTP1-GFP / WT heterografts (Fig. 2A-E), whether in the agar-grown seedlings (Fig. 2 A-C, 8 to 13 DAG) or the older soil-grown plants (Fig. 2E, 80 DAG). Imaging the scion with the same microscope settings completely saturated the confocal photomultiplier (images therefore not shown). GFP fluorescence localised along the vascular strands, with acropetally increasing intensity towards the root tip, down to about 250 µm from the quiescent centre (251 ± 11.8 µm at 13 DAG, n = 5), within the transition zone from the root elongation zone to the root meristem, where the GFP signal completely disappeared (Fig. 2A, B, E; Supplementary Fig. 1), coinciding with the end of the protophloem. Closer inspection at higher resolution along the root elongation and differentiation zones showed patchy GFP signal intensity reflecting preferential protein accumulation in phloem-pole pericycle cells at the sites of lateral root initiation (Fig. 2C, F, G). WT roots grafted to scions expressing a p35S∷YFP-cTCTP1 construct providing a much stronger fluorescence signal, also clearly showed pericycle-specific localisation of YFP fluorescence (Fig. 2D). TCTP1-GFP fluorescence was also detected in LR primordia (LRP), but was much weaker, sometimes barely detectable until the root had emerged and started to fast elongate (Fig. 2H). The pattern of GFP fluorescence in control roots grafted on a scion expressing GFP alone under the same promoter was very different, with a very high ubiquitous signal in the whole stele and throughout the root meristem (Supplementary Fig. 2), as previously reported [51, 52]. Taken together, these results suggest that rootward *TCTP1* mobility is actively controlled and may have a specific signalling function in root development, targeted to cells involved in the spatial patterning of lateral root primordia along the primary root.

**Fig. 2.**
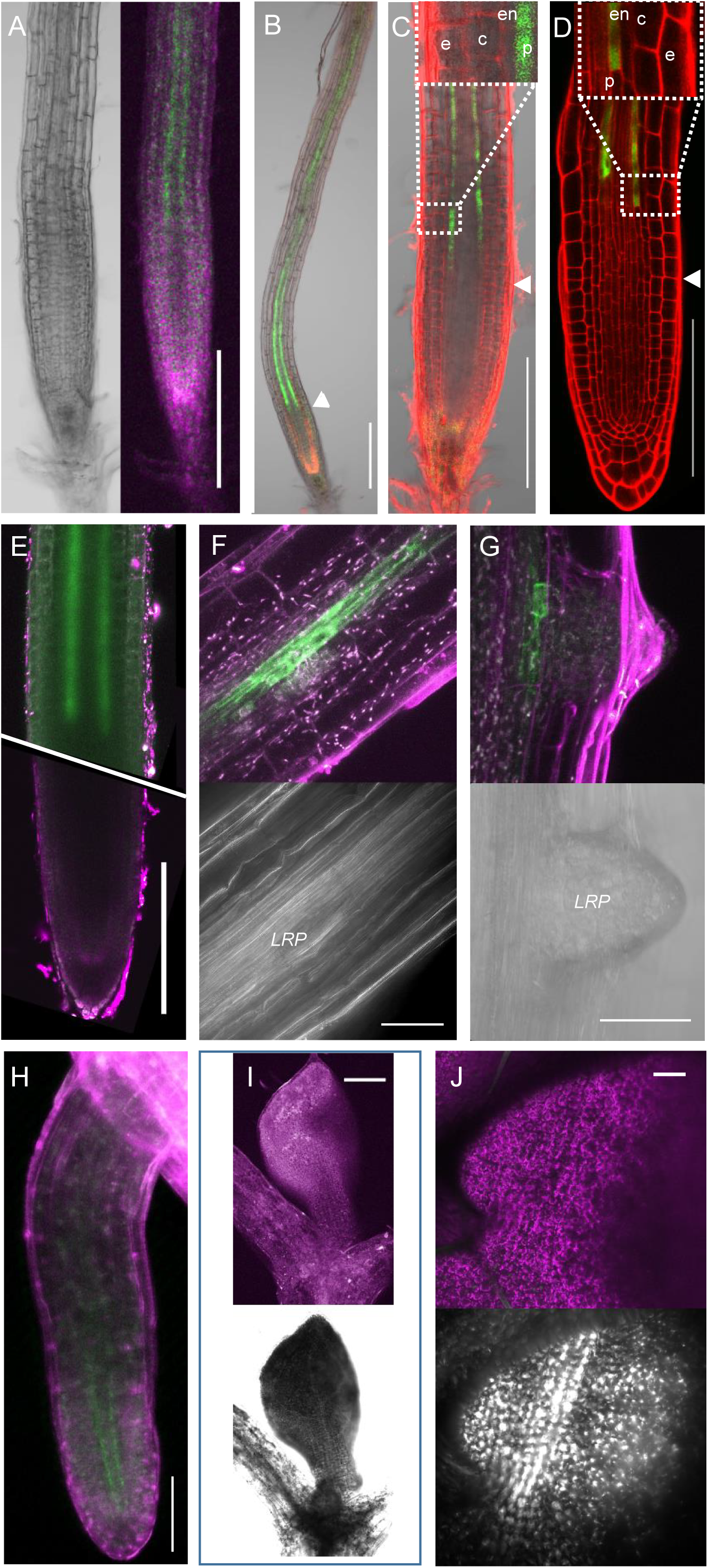
TCTP1-GFP distribution in grafted WT rootstock is restricted to the pericycle, with preferential accumulation in phloem-pericycle cells at sites of lateral root formation. **A-H** Confocal laser microscopy images of WT roots grafted onto scions expressing *pTCTP1∷gTCTP1-GFP* (**A-C** and **E-H**) or *p35S∷cYFP-TCTP1* (**D**). Roots were imaged in seedlings grown in agar plates, 8 DAG (**A**) or 13 DAG (**B-D**) (n ≥ 30 primary roots), and in flowering soil-grown plants, 80 DAG (**E-H**), n ≥ 10; primary root and *LRP* (**E-G**); elon-gating lateral root (**H**). Note the localisation of the GFP fluores-cence in parallel strands, the increasing signal closest to the root tip and the preferential accu-mulation of the GFP-tagged TCTP1 protein in phloem-pole pericycle cells (**C-D** and **F-G**) at the sites of lateral root initiation (label *LRP* in **F-G**). GFP fluores-cence (green) and auto-fluores-cence (magenta/red) were sepa-rated by spectral unmixing. Inserts in **C** and **D** show TCTP1-GFP (**C**) and YFP-TCTP1 (**D**) localisation, respectively, in the pericycle (p); GFP fluores-cence was undetectable in the epidermis (e), the endodermis (en), or the cortex (c). **I-J**, Absence of detectable GFP fluorescence signal in WT scions grafted to *pTCTP1∷gTCTP1-GFP* roots, 8 DAG in agar-grown plants (**I**) or 80 DAG in soil-grown

To examine this in more detail, we monitored the appearance of TCTP1-GFP fluorescence in the rootstock of TCTP1*-*GFP / WT grafts over a ten day period following grafting. The earliest evidence of GFP fluorescence was on 7 DAG (23 out of 28 roots; Supplementary Fig. 3), consistent with reports of fully functional graft junction in Arabidopsis [53]. Strikingly, the preferential TCTP1-GFP accumulation in the primary root elongation zone observed in older roots (Fig. 2), was already obvious (Fig. 3A). Moreover, when imaging the entire root from base to tip, TCTP1-GFP fluorescence was first encountered in two patches localised in pericycle cells at the base of the two youngest LR primordia, both at initiation stages I-II [3] (Fig. 3H-I). This result further supports the notion of a destination-selective signalling function of mobile *TCTP1* gene products originating from the shoot, in the initiation of lateral roots. Given the bi-directional mobility of *TCTP1* mRNA (Fig. 1G, H) we examined the presence of GFP-fluorescence in WT scions grafted onto *pAtTCTP1∷gAtTCTP1-GFP* roots. GFP fluorescence was undetectable, whether in young or mature leaves, or in the shoot apical meristem (Fig. 2I, J).

**Fig. 3.**
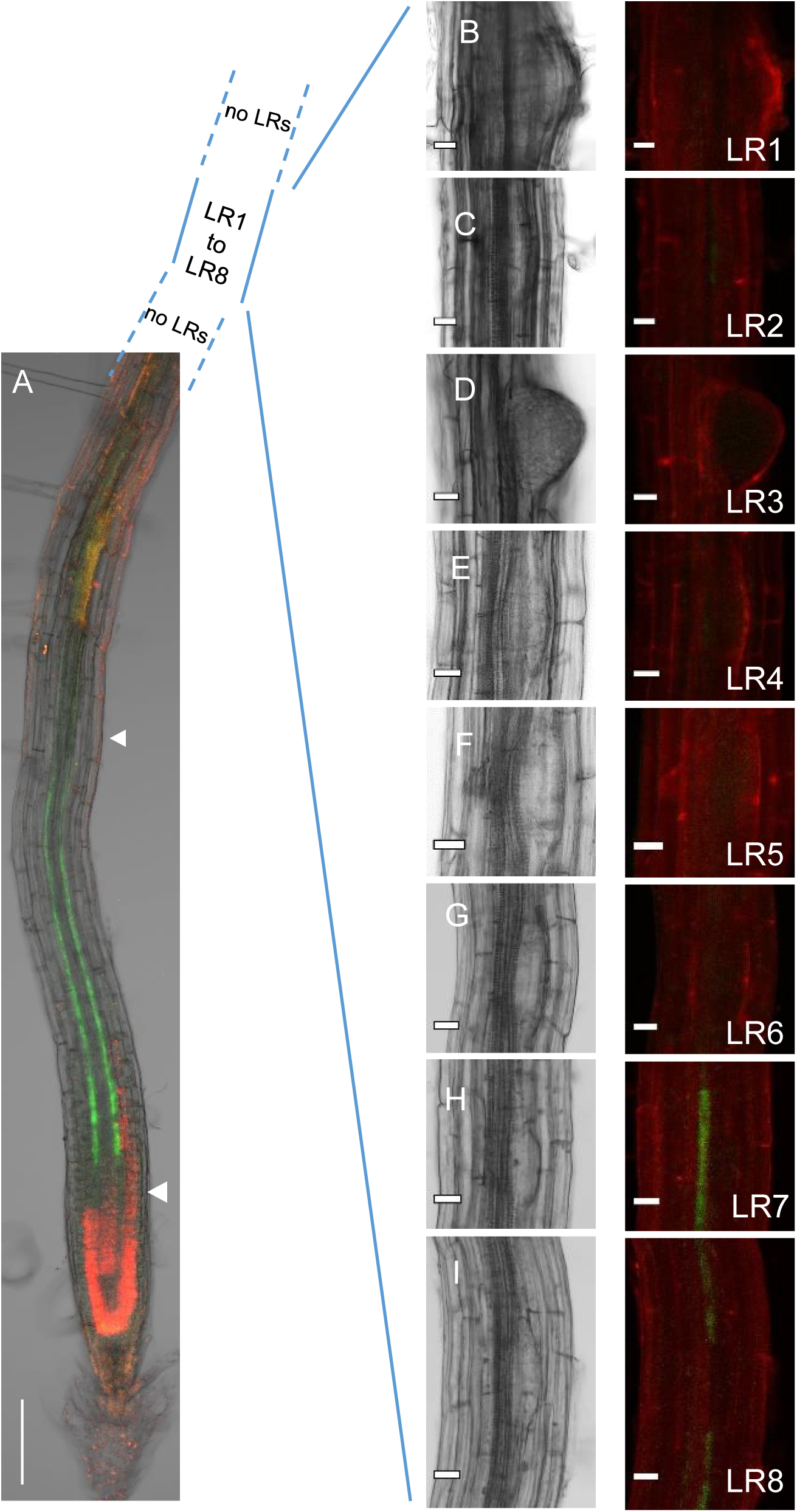
Scion-derived TCTP1-GFP fluorescence in grafted WT rootstocks is first detected at the most apical lateral root initiation sites. **A,** Confocal laser scanning microscopy image of a WT primary root grafted to a scion expressing *pTCTP1∷gTCTP1-GFP*, 7 DAG. Scale bar, 200 µm. **B** to **I**, CLSM images of lateral root primordia along the primary root, from the most distal (**B**) to the most apical primordium (**I**) closest to the root tip, just above the zone of *LRP* priming. For each primordium, transmission channel on the left, fluorescence channel on the right.TCTP1-GFP fluorescence (green) is only visible at the initiation sites of the two youngest *LRP* primordia (**M** and **N**). Scale bars, 20 µm.

### Constitutive expression of *AtTCTP1* in scion promotes scion growth which in turn stimulates root growth

In our earlier characterisation of *AtTCTP1* we showed that *AtTCTP1* gene products, mRNA and protein, are constitutively highly expressed in the primary root meristem and LR primordia, and that *AtTCTP1* silencing inhibits root elongation and lateral root branching [35]. The above results in the present study suggested a role of AtTCTP1 in root development through mobility too. To investigate that, for lack of roots devoid of constitutive *AtTCTP1* expression given the embryo lethality of total *AtTCTP1* knock-out and the dwarfism of *tctp1* seedlings rescued through embryo culture [35, 36], we severed TCTP1-RNAi roots from 5 days old seedlings expressing a constitutive *AtTCTP1* silencing construct [35], (thereafter referred to as TCTP1-RNAi or RNAi) and grafted them to either a WT scion of the same age, or back to the severed homologous TCTP1-RNAi scion. Root development in these grafts was then monitored over the next 3-4 weeks. We reasoned that the 100-fold higher *AtTCTP1* constitutive expression in WT scions than TCTP1-RNAi scions [35] should translate into a significantly increased amount of scion-to-root mobile *AtTCTP1* messenger and protein in WT / RNAi compared to RNAi / RNAi grafts, and thus enable us to determine whether the distinctive short root and reduced branching of the rootstock is root-autonomous or involves signalling by *AtTCTP1* from the scion. WT / RNAi reciprocal heterografts were grown alongside each other and control WT/ WT and RNAi / RNAi homografts, in replicated plates. WT primary roots grew faster than TCTP1-RNAi roots irrespective of scion genotype (Fig. 4A, B). Both WT and TCTP1-RNAi roots elongated faster when grafted onto a WT rather than TCTP1-RNAi scion (26% and 21%,increase in maximum relative elongation rate 13 DAG, respectively, Fig. 4C). It is known, however, that siRNA can move long distances through the plant [54–57]. To test whether this explained the root growth inhibition associated with TCTP1-RNAi scions, we measured *AtTCTP1* transcript abundance in homo- and hetero-grafts scions and rootstocks by quantitative RT-PCR. *AtTCTP1* mRNA levels in WT roots showed a nearly 6-fold reduction in RNAi / WT compared to WT / WT seedlings (*P = 0*.*005,* Fig. 4D). This is much higher than could be expected from simply a reduction of mobile rootward *AtTCTP1* mRNAs (see Fig. 1G,H) and hence suggested some down-regulation of *AtTCTP1* expression in these roots by mobile siRNA from the RNAi scion. By contrast, *AtTCTP1* transcript abundance in RNAi roots was similar regardless of scion genotype (0.68± 0.118 and 0.77± 0.013 relative transcript abundance in RNAi / RNAi and WT / RNAi grafts, respectively, *P* = 0.55; Fig. 4D), ruling out that simple explanation for their slower elongation rate. To verify that mobile *AtTCTP1* transcripts from the scion were not silenced by siRNA upon delivery to the root, we grafted TCTP1-GFP scions on TCTP1-RNAi roots and quantified transgenic *TCTP1-GFP* mRNAs in the root stock (Fig. 4E). *TCTP1-GFP* messengers were present in roots of TCTP1-GFP / RNAi grafts, in low but significant abundance representing a similar or higher fraction of the amount of TCTP1 mRNAs transcribed in the scion of TCTP1-GFP / WT grafts in our earlier experiments (ca 0.006 %). Moreover, GFP fluorescence from the encoded TCTP1-GFP protein was consistently detected in the root, with the same spatial expression pattern (Fig 4F compared to Fig. 2B). Altogether these data indicate the presence of a significantly higher amount of intact *TCTP1* messengers of scion origin in TCTP1-RNAi roots grafted to a WT scion instead of homologous TCTP1-RNAi scion.

**Fig. 4.**
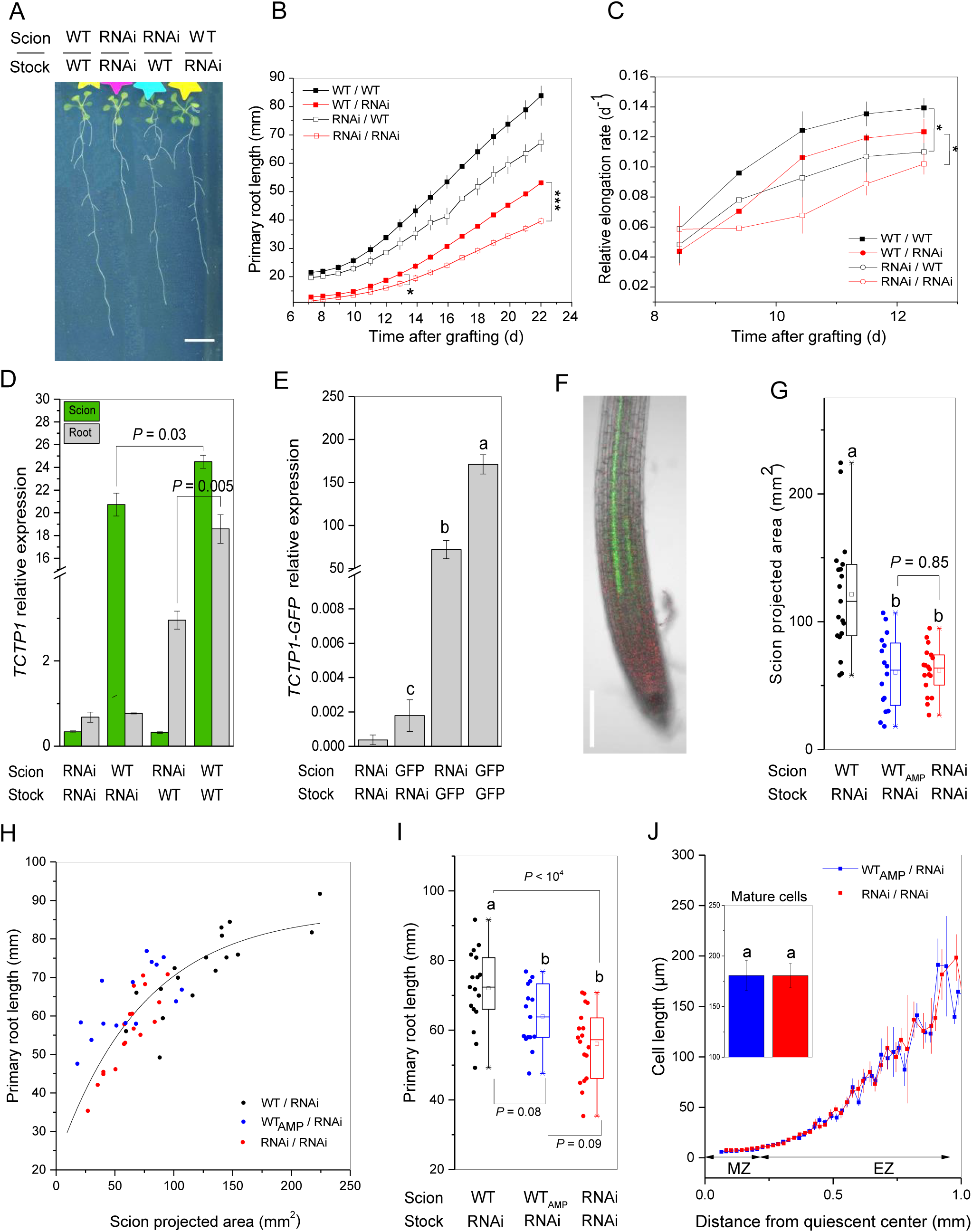
Constitutive expression of *AtTCTP1* in scion promotes scion growth which in turn stimulates primary root elongation. **A,** Representative images of reciprocal grafts between WT and *TCTP1*-RNAi lines, and control WT / WT and TCTP1-RNAi / TCTP1-RNAi homografts, 22 DAG. Scale bar, 10 mm. **B** and **C**, Primary root lengths (**B**) and relative root elongation rates (**C**) *versus* time. Means ± SE, n = 6-9. Asterisks in **B** denote statistically significant differences by two tails Student’s T-test; * *P*< 0.05; *** *P*< 0.001. **D**, *AtTCTP1* relative gene expression levels in rootstocks and scions, 22 DAG (means ± SE, n = 3 pools of at least 5 plants each). Probability levels (*P*) of *s*tatistical significance was determined by two tails Student’s T-test. **E**, *AtTCTP1-GFP* relative expression in the rootstock of reciprocal grafts between TCTP1-RNAi and TCTP1-GFP lines (denoted for brevity “RNAi” and “GFP”, respectively, in the Figure), and control homografts. Different letters indicate statistically significant differences, above noise levels in roots from RNAi/RNAi grafts (one-way ANOVA and Tuckey’s test, n= 4 pools of ≥ 6 roots). **F**, Representative image of GFP fluorescence in TCTP1-RNAi roots grafted to TCTP1-GFP scions. Scale bar, 200 µm. **G** to **J**, Comparison of grafted seedlings sharing the same rootstock (TCTP1-RNAi) but differing in scion genotype or size (“AMP” subscript denotes WT scions trimmed 7 DAG, see Methods): **G,** scion sizes 18 DAG; **H,** individual primary root lengths *versus* scion projected area; **I,** primary root lengths. **G** to **I,** Dots represent individual seedlings, n = 16-19 seedlings per graft type; **G**-**I,** boxes show median, first and third quartiles, and upper and lower whiskers give a graphic representation of the interval containing all data points within ± [1.5×(Q_3_-Q_1_)] range. **J**, Epidermal cell lengths along the primary root meristem (MZ) and elongation zone (EZ), n = 5 roots per graft type. Shown are moving averages of cellular lengths over 20µm windows. The inset depicts mature cell lengths (means ± SE). Different letters in **G, I** and **J** denote statistically significant differences by Student’s T-test (*P* < 0.05 unless indicated).

Despite being all trimmed to a similar size at the time of grafting (see Methods), WT scions quickly became larger than TCTP1-RNAi scions (Supplementary Fig. 4), consistent with the high expression of *AtTCTP1* in the shoot apical meristem and its growth promoting effect in leaves [35]. This suggested that the faster elongation rates of TCTP1-RNAi roots in WT / RNAi than RNAi / RNAi grafts might then at least partly reflect an increased photo-assimilate supply from larger scions, rather than higher abundance of *AtTCTP1* mRNA translocated from the scion. To address this, we grafted TCTP1-RNAI roots onto WT scions or onto homologous TCTP1-RNAi scions of the same age as earlier. But at 7 DAG, when the graft junction was fully established and TCTP1-RNAi root lengths were still similar regardless of scion genotype (12.8 ± 1.25 mm and 11.4 ± 0.91 mm in WT / RNAi and RNAi / RNAi grafts, respectively;Student’s T-test, *P* = 0.13, n ≥ 6, Fig. 4B), we normalised WT scion sizes to RNAi scion sizes through amputation of one cotyledon (scions thereafter denoted WT_AMP_). To minimise potential confounding wounding effects in subsequent phenotypic analyses, the petiole of one cotyledon was manually pinched in control grafts. Two weeks later (22 DAG), scion sizes were still similar in the two sets of grafts (WT_AMP_ / RNAi and RNAi / RNAi, Fig. 4G, Student’s T-test, *P* = 0.85, n ≥ 16), and about half the size of non-amputated WT scions in WT / RNAi grafts. Remarkably, associated TCTP1-RNAi rootstock showed similar primary root lengths (Fig. 4H), significantly shorter than roots of WT / RNAi grafts. This was confirmed in independent experiments (Supplementary Fig. 5). When individually plotted against scion sizes, root lengths described a unique relationship for the three sets of grafts, and data points for TCTP1-RNAi roots associated to WT_AMP_ or TCTP1-RNAi scions overlapped (Fig. 4H). Consistently, relative root elongation rates over the monitoring period (7 to 22 DAG) were similar (4.17 ± 0.15 h^-1^ and 4.07 ± 0.14 h^-1^ in WT_AMP_ / RNAi and RNAi / RNAi grafts, respectively, Student’s T-test, *P* = 0.09, n ≥ 16). Moreover, the cell length profiles in the root elongation zone also showed complete overlap, and final cell lengths were similar (Fig 4J). These results indicate that, at same scion size, differences in constitutive *AtTCTP1* expression levels in the scion and rootward mobile *AtTCTP1* gene products have little impact on primary root elongation.

**Fig. 5.**
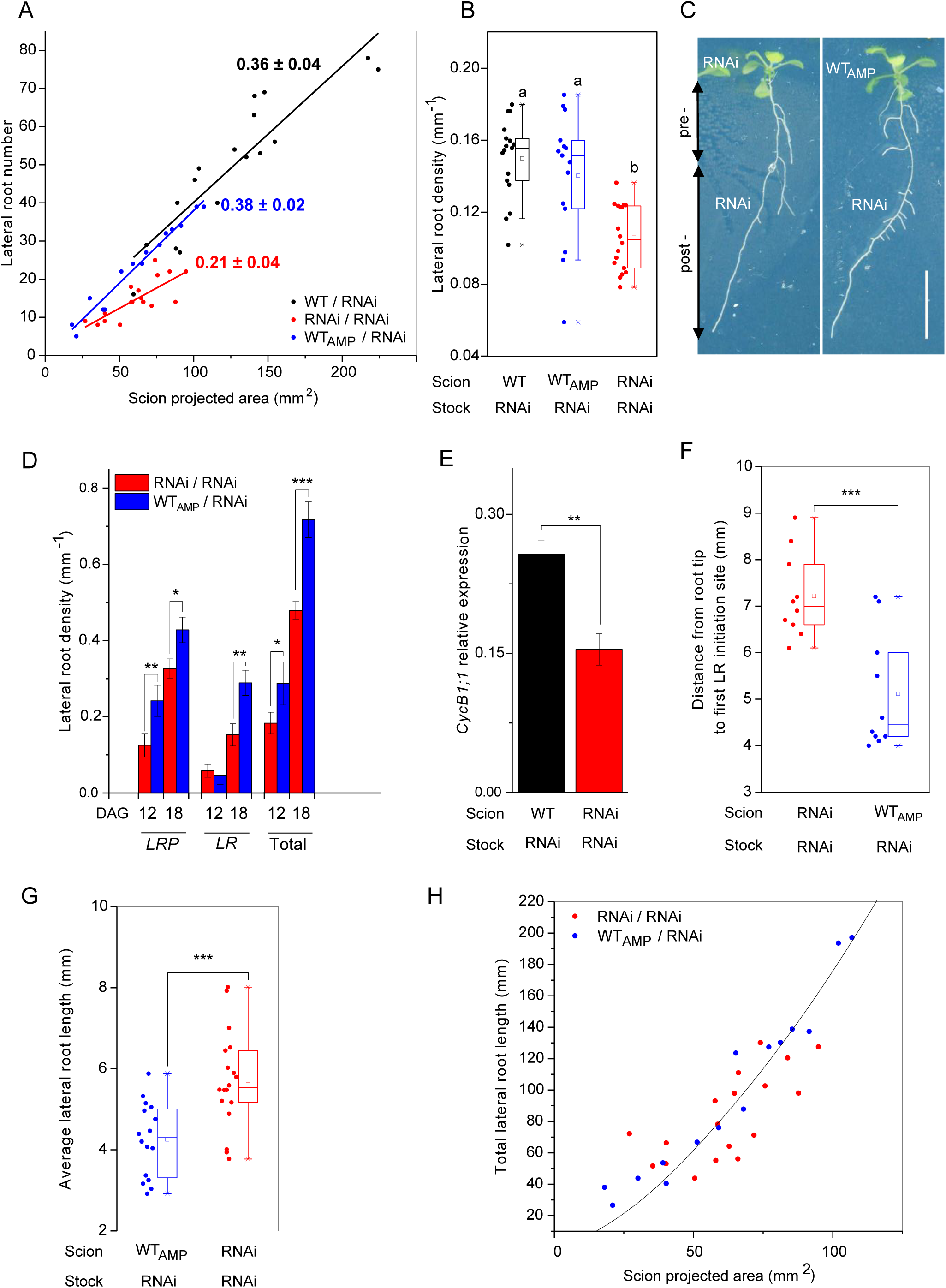
Rootward TCTP1 movement promotes lateral root initiation and emergence. **A**, Lateral root number as a function of scion projected area; linear regression lines and slopes ± SE are shown for each set of grafts, n = 15-17. **B**, Lateral root density (number of lateral roots per unit length of primary root). Different letters indicate statistically significant differences by one-way ANOVA followed by Bonferroni posthoc test, n = 15-17 roots. **C**, Representative photographs of a *TCTP1*-RNAi / *TCTP1*-RNAi homograft (left) and a WT_AMP_ / *TCTP1*-RNAi heterograft (right), 18 DAG: “pre-” and “post” denote the portion of primary root formed pre- and post-grafting and scion size normalization (0 mm −8 mm and 10 mm to root tip, respectively). Scale bar, 10 mm. **D**, Lateral root density on the primary root portion formed after scion size normalization. *LRP* and *LR* denote non-emerged Lateral Root Primordia and elongating Lateral Roots, respectively. Means ± SE, n = 14 roots. **E**, *Cyclin B1;1* relative expression measured by qRT-PCR in rootstocks sampled 10 DAG. Means ± SE, n = 4 biological replicates, each consisting of at least 6 pooled rootstocks. **F**, Distance between the youngest, most proximal LRP and the root tip, n=10. **G**, Average lateral root length, and (**H**) total lateral root length as a function of scion projected area, measured 18 DAG. **D** to **G**, Statistical significance was determined by two tails Student’s T-test (* *P* <0.05; ** P < 0.01; *** *P* < 0.001, n=14-18). **A-B**, and **G-H**, Each data point represents an individual plant.

### Graft-mobile *AtTCTP1* transcripts and encoded protein promote lateral root initiation and emergence

As scion-derived TCTP1-GFP showed preferential accumulation at sites of lateral root initiation (Fig. 2), we next closely examined root branching patterns. The number of lateral roots varied between plants. That variation was closely correlated to variation in scion size (Fig. 5A). Data points for WT_AMP_ / RNAi and WT / RNAi grafts fell on the same line (slopes 0.38 ± 0.02 and 0.36 ± 0.04, respectively), indicating that partial amputation of WT_AMP_ scions had *per se* no unwanted confounding effects on root development. Remarkably, lateral root numbers in RNAi / RNAi grafts fell significantly below those seen in WT_AMP_ / RNAi heterografts. The density of lateral root formation sites along the primary root was decreased by 41% on average (Fig. 5B), while being as high in roots grafted to WT_AMP_ than the much larger WT scion (Student’s T-test, *P* = 0.36, n ≥ 15; Fig. 5B).

We next used Differential Interference Contrast (DIC) microscopy to examine the entire length of the primary root in WT_AMP_ / RNAi and RNAi / RNAi grafts, for non-emerged LR primordia (denoted *LRP* hereafter; stages I-VII [3]) in addition to emerged LRs (denoted *LR*), also separately scoring those found on the primary root segment formed pre-or post-grafting and scion-size normalisation (Fig. 5C). The former were confined to the basal 8-9 mm of the primary root, and in total amounted to 2.9 average in the two sets of grafts (Student’s T-test, *P*= 0.89, n = 14). The density of both *LRP* and *LR* lateral roots on the younger, proximal primary root segment formed post-scion size normalisation, was significantly greater in WT_AMP_ / RNAi than RNAi / RNAi grafts (*P* < 0.05, n = 14; Fig. 5C, D). This was already clear 12 DAG and even more obvious 6 days later. Consistent with the increased *LRP* density, the expression of *CycB1;1,* a marker of the early divisions of pericycle cells that initiate lateral root formation [58] was also increased in roots from WT_AMP_ / RNAi grafts compared to those of RNAi / RNAi grafts (*P* <0.004, Fig. 5E). In the course of these experiments, we noticed that the youngest, most acropetal *LRP* seemed located closer to the root tip. Systematic measurements of its coordinate along the primary root showed that the zone of lateral root formation indeed extended significantly closer to the root meristem in WT_AMP_ / RNAi than RNAi / RNAi grafts (*P* = 4.10^-4^, n = 10, Fig. 5F). Root patterning was completely normal (Supplementary Fig. 6). Taken together, these data indicate a stimulation of lateral root initiation and emergence in WT_AMP_ / RNAi grafts, at same scion size and primary root length and anatomy, thus likely related to enhanced delivery of mobile *AtTCTP1* gene product(s) from WT scions compared to RNAi scions. The identical cell length profiles along the whole transition, elongation, and differentiation zones of the two sets of roots imply an increased frequency of LR priming events along the primary root pericycle.

**Fig. 6.**
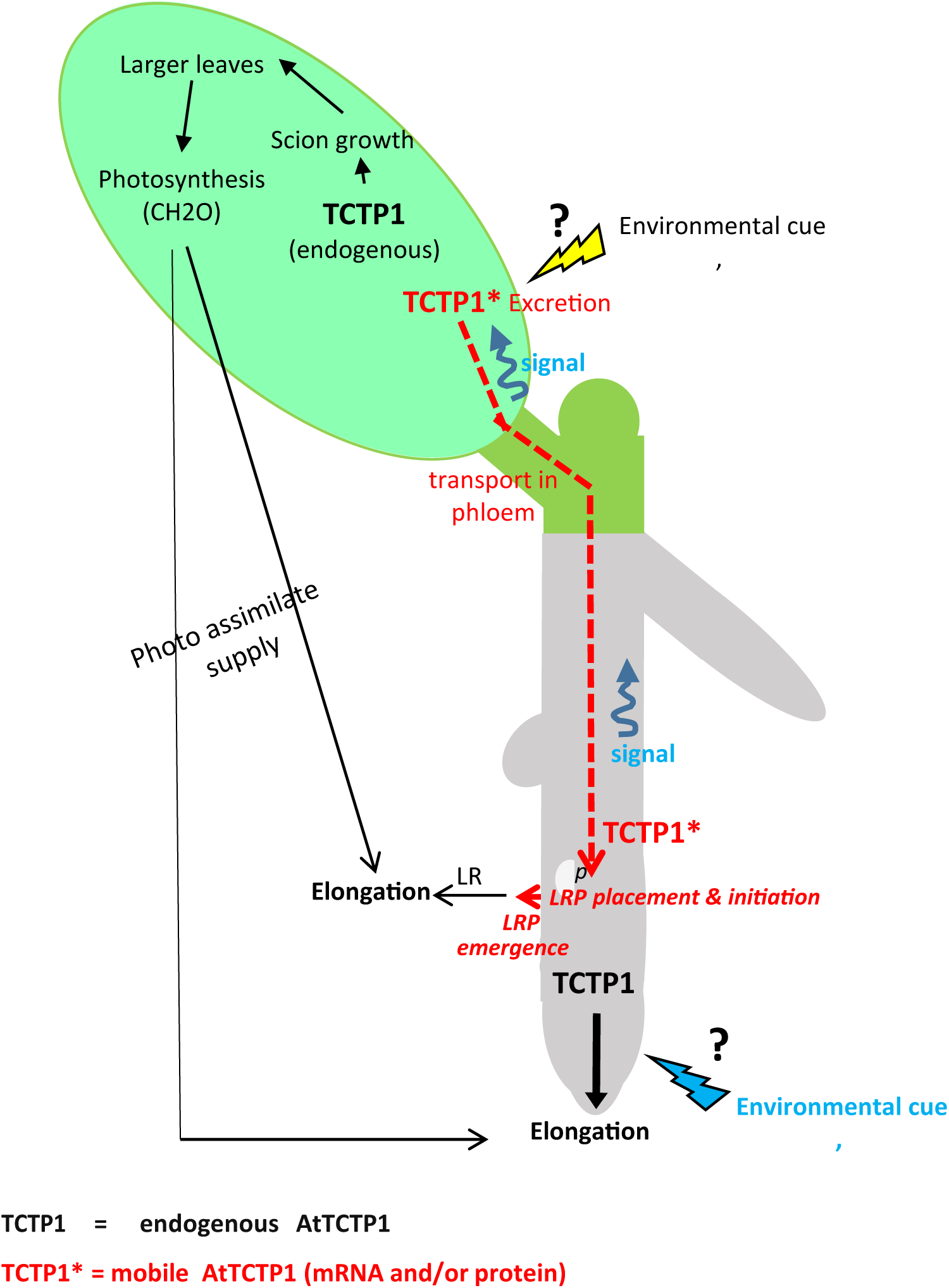
Proposed model of synergistic regulation of root development by constitutive and mobile *AtTCTP1* gene products. Local constitutive At*TCTP1* (black lettering and arrows) acts as a positive regulator of cell growth and prolifera-tion within roots and aerial organs and thus overall organ size. Overall root length is bounded by photo assimi-late supply, hence indirectly dependent on AtTCTP1 constitutive expression in the shoot through its effects on the size of the photosynthetic apparatus. Mobile *AtTCTP1* messengers translocated from the shoot and encod-ed proteins (red lettering and arrows) act as systemic destination-selective signalling molecules to dynamically modulate the spatio-temporal patterning of lateral root initiation and emergence sites on the primary root, and the size of the LR formation zone, perhaps too the initial “priming” step of pericycle *LRP* founder cells. The mechanisms flagging *AtTCTP1* gene product(s) for excretion and export to roots, and unloading in destination cells are unknown, but likely regulated by a combination of above-ground and below-ground environmental cues in interaction with endogenous cues, including age-dependent. Dashed arrows depict “movement”; full arrows denote “promotion”. *p* denotes the pericycle.

While initiated and emerging in greater number in WT_AMP_ / RNAi than RNAi / RNAi grafts, lateral roots were on average shorter (*P* = 4.10^-4^, n ≥ 16; Fig. 5G). Strikingly, the overall result was that their cumulated length over the whole primary root was unaffected, similar to that measured for RNAi / RNAi grafts (*P* = 0.32, n ≥ 15, Fig. 5H; Supplementary Fig. 7). Plant to plant variation in cumulated LR root length was correlated to individual variation in scion size (Fig. 5H) as found for primary root length, and data points fell on the same relationship for the two sets of grafts (Fig. 5H). Taken as a whole, these results indicate a signalling role of *AtTCTP1* rootward mobility in root development, specifically targeted at the regulation of early events of LR formation and of LRP emergence from covering layers.

## Discussion

TCTP function and physiological roles in plants are only beginning to be unravelled. The possibility of a dual role, as a protein acting not only cell-autonomously, but also extra-cellularly as found in mammals and other eukaryotes remains unknown. This study indicates a targeted, selective function of scion-encoded mobile endogenous Arabidopsis *AtTCTP1* mRNA and translated protein in shaping the deployment of the root system.

*AtTCTP1* mRNA movement was reported to occur in a strictly rootward direction [11]. Here, we consistently detected a bi-directional movement, whether in plate assays with young seedlings raised under similar conditions as in the earlier study, or after graft transfer to soil. This discrepancy may be related to the different reporter constructs used: full genomic *TCTP1* here, including 5’UTR and native upstream promoter region; *TCTP1* cDNA fused to the constitutive cauliflower mosaic virus promoter in the previous study. The 5’ UTR of *AtTCTP1* contains a conserved 5’TOP (terminal oligopyrimidine tract) motif, and two AUUUA motifs are present in the 3’UTR. TOP mRNAs are highly sensitive to translational control and AUUUA motifs are also important in mRNA stability and translation [59]. In animals these motifs are thought to be important in the control of TCTP translation [33] and in Arabidopsis itself have been suggested to influence the expression pattern of AtTCTP1 [36]. It may be that the 5’ and 3’UTRs also play a role in the mobility and/or transport of *AtTCTP1* transcripts, as shown for *StBEL5* mRNA movement from leaf and petioles to stolon tips in the regulation of tuber formation in *Solanum tuberosum* [27]. Supporting our results, an also bi-directional endogenous movement of the native *Vitis vinifera TCTP1* mRNA homologous to *AtTCTP1* has been documented in heterografts of two polymorphic *Vitis* genotypes, through genome-wide sequencing of scions and rootstock mRNAs [13]. The consistency of our observations in seedlings grown in vitro and older soil-grown plants provides evidence that long distance movement of endogenous *AtTCTP1* mRNA between shoot and root is not a transient occurrence, nor an experimental artefact, but a sustained phenomenon, which takes place under physiological conditions.

If that phenomenon has a physiological function one would expect it to be actively controlled, in a growth conditions or development stage dependent manner. Fitting with this, while always representing a small fraction of the transcripts produced in the source organ as seems the norm [12, 13, 22], the proportion of At*TCTP1* mRNAs transmitted across root-shoot graft junctions showed significant variation from plant to plant and between *in vitro* and soil experiments (Fig. 1). Consistently, for rootward movement, the fluorescence signal associated with TCTP1-GFP protein derived from mobile scion *TCTP1 mRNAs* was also of variable intensity, especially in the population of elongating lateral roots of soil-grown plants, being even undetectable in some. No conclusion is possible about TCTP1-GFP protein in the scion of reciprocal WT / TCTP1-GFP grafts. It was undetectable, indicating either absence of translation at least under our experimental conditions, or very low abundance but nevertheless functionality as shown for some other regulatory proteins of root development such as BREVIS RADIX for example [60].

Another expectation of mobile gene products serving a physiological function, is that they give rise to distinct, quantifiable phenotypes. To examine this we focused on *AtTCTP1* rootward mobility. Our results provide convergent evidence towards a systemic signalling role targeted at cells involved in the initiation and emergence of lateral roots. First, the scion-derived TCTP-GFP protein detected in the WT primary root of TCTP1-GFP/WT grafts consistently showed a distinctive localisation pattern, being a) absent from the root meristem, b) confined to the vasculature, with recurrent peaks of higher intensity, systematically coinciding with sites of lateral root primordia initiation in the pericycle; c) highly abundant in the root elongation and transition zones upstream to the root meristem proper, which encompass a region of “oscillatory gene expression” where priming of LR initiation takes place [4, 5]. This pattern is in stark contrast with the ubiquitous expression, in all root layers and also the root meristem, of the constitutive root AtTCTP1 protein, translated from locally transcribed *AtTCTP1* transcripts [35]. It is also distinct from the pattern observed for mobile GFP or YFP alone, and passive mass flow transport through the phloem and leakage from companion cells to pericycle cells. These results indicate the presence of active targeting, capture or unloading mechanisms in the root of *AtTCTP1* gene products encoded in the shoot. Second, TCTP1-RNAi roots of similar structure, anatomy, size and elongation rate but associated with WT_AMP_ scions rather than TCTP1-RNAi scions of similar size, exhibited an extended zone of lateral root formation starting closer to the root tip. Third, the densities of root branching sites (emerged LR) and of lateral root initiation sites (non-emerged LRP at more acropetal positions) along that zone were both increased. The specificity of these changes and shift towards more numerous and on average shorter LR, in an extended zone, while the overall length of root over the whole root system was unaffected, contrasts with the general promotion of both primary and lateral root elongation associated with increased local constitutive AtTCTP1 expression [35], or with increased scion size (Fig.4 and 5), or with independently induced increases of photo assimilate supply via stimulation of carbon metabolism [61, 62]. Differential sucrose uptake by TCTP1-RNAi and WT_AMP_ scions directly from the media – which is known to influence lateral root emergence [63] - can also be ruled out as an explanatory factor for the effects of scion genotype observed here on root system architecture, as our grafts were raised in the absence of exogenous sucrose supply.

The phloem is the long distance delivery path to roots of auxin synthesised in leaves and cotyledons. Auxin has a pivotal role in orchestrating root architecture [64–66], being required for the activation of LR potential initiation sites in the pericycle, the subsequent process of primordium initiation and formation, and the breaking of overlying tissues for its emergence out of the primary root epidermis [1, 4, 67–72]. However, TCTP1-RNAi seedlings of the same age as used in this study and raised under similar growth conditions in earlier experiments, were found to in fact have higher endogenous auxin concentrations than WT seedlings in both shoots and roots [35], and, consistently, higher expression of the auxin inducible transcriptional regulator IAA5. In addition, application of the auxin polar transport inhibitor NPA in the present study had no detectable influence on the abundance or localisation of the AtTCTP1 protein translated from mobile *AtTCTP1* transcripts (Supplementary Fig. 8). It is therefore unlikely that stimulation of lateral root initiation and emergence in WT_AMP_ / RNAi compared to RNAi / RNAi grafts could result from a higher auxin production in the scion and increased auxin delivery to the root.

Taken as a whole, these observations provide a compelling argument for ascribing the scion genotype-dependent modulation of root architecture observed in our grafts to differential transmission rate of mobile scion *AtTCTP1* gene products to the root, associated with the large (two orders of magnitude) difference in constitutive *AtTCTP1* transcripts abundance between WT_*AMP*_ and TCTP1-RNAi scions (Fig. 4D [35]). We propose a model (Fig. 6) where local constitutive At*TCTP1* controls core cellular processes, vital to cellular function, growth and proliferation, in both roots and shoot, and determines the overall length of root that can be formed in interaction with photo assimilate supply; while mobile *AtTCTP1* messengers -and perhaps proteins-transmitted from the shoot act as destination-selective systemic signalling molecules to dynamically modulate the spatio-temporal pattern of lateral root initiation and emergence, and possibly too the initial priming of pericycle LRP founder cells, i.e. the plasticity of root system architecture.

Whether *AtTCTP1* mRNAs only get transmitted from shoot to roots, and fulfil their function through destination-selective delivery and decoding in the root, or whether some might get translated in the scion, followed by selective loading of the protein into the phloem, and selective unloading in specific destination cells in the root, remains to be determined. The two scenarios are not exclusive. Disentangling them will be challenging given the high and ubiquitous constitutive expression of AtTCTP1 in source and destination tissues of mobile *AtTCTP1* mRNAs, and also the limitations of transgenic approaches with non-endogenous gene products when it comes to characterisation of movement and function. Suggesting the possibility of AtTCTP1 protein mobility too, an actively controlled rootward movement of the highly similar pumpkin CmTCTP1 (CmaCh11G012000) in rice phloem sieve tubes has been reported[50]. Its functionality was not analysed, but most interestingly the mobile CmTCTP1 (CmaCh11G012000) was found to be translocated as part of a protein complex including CmPP16-1 and CmPP16-2 phloem RNA binding proteins and also the initiation factor CmeIF5A, among other undetermined components -an association fitting with the assumed molecular function of plant TCTPs in protein synthesis, as established in animals [73, 74]. Under either scenario – mobility of the messenger only, or of the protein also-, it will be intriguing to elucidate how destination cells recognise the mobile *AtTCTP1* gene products generated in distant cells from those they autonomously produce; and what their mechanisms of action are.

The plasticity of root system architecture is pivotal to plant adaptation to changes in environmental conditions, and strategic mining of spatially and temporally variable soil resources. *AtTCTP1* appears as a central controller of that plasticity. Combining a dual function - general constitutive growth promoter, and mobile signalling agent between shoot and roots - in a vital, highly expressed gene, furthermore encoding a protein highly sensitive to translational and post-translational modifications, appears as a clever strategy for a sensitive and dynamic tuning of root system architecture to optimise the compromise for roots between reaching deeper or branching more profusely according to growth. Interestingly, the much lowlier and more specifically expressed Arabidopsis *AtTCTP1* orthologue, *AtTCTP2*, was reported to encode a mRNA and protein with bi-directional long distance mobility when overexpressed in *Nicotania benthiamana* [29]. And movement across the graft junction appeared to correlate with the formation of aerial roots at the interface of the grafted stem segments [29]. This raises the possibility of a broad role of *TCTP* mobile mRNAs/proteins in *de novo* root organogenesis, whether lateral roots or shoot-born roots, with specificity between gene isoforms in cell types targeted for developmental reprogramming and formation of root primordia from a variety of tissues [2], as most appropriate.

Plant TCTP genes show high similarity among species. TCTP messengers and proteins have been detected in the vasculature of diverse species, including the monocot rice where the TCTP family is reduced to one member (Supplementary Table 1). This suggests that the mobility and extracellular signalling function of *AtTCTP1* to control root organogenesis might be widely conserved within the plant kingdom, and highly relevant to a better understanding of post-embryonic formation of lateral organs in plants, and the elusive coordination of shoot and root morphogenesis.

## Methods

### Plant material and Growth conditions

The transgenic proTCTP1∷gTCTP1-GFP∷NOS (TCTP1-GFP) and the TCTP1-RNAi lines are as described in our previous work [35], and the 35S∷YFP-cDNATCTP1 as in [11]. All lines are in Arabidopsis thaliana Columbia (Col-0) background (wild type, WT). Seeds used within individual experiments are of the same age and were harvested from isolated plants grown together, under the same conditions.

The hypocotyl micro-grafting method was as described [75] with minor adaptations to improve success rate rate and reduce adventitious root emergence. Seeds were surface-sterilized for 5 min with a solution of sodium hypochlorite 0.125% (v/v) and 90% (v/v) ethanol, rinsed 3 times in 100% ethanol and dried under a laminar flow hood, before being resuspended in sterile water and sown directly on the surface of a nitrocellulose membrane strip (Membrane Filter – Cellulose Nitrate 0.45 µm diameter, Whatman code NC45ST) laid on top of a nutrient agar gel (2,15 g L^-1^ MS salts, 0.5% sucrose, 1% Agar type-M, pH 5.7) in petri plates. The plates were sealed with porous micropore tape (3M), stratified for two days at 4°C in the dark and incubated under controlled conditions (12h photoperiod, 21 °C, 120 µmol quanta m^-2^ s light intensity), in a vertical position. Four to six days later, under sterile conditions, the two cotyledons were severed mid-way through their petiole. The seedlings were positioned with the hypocotyl perpendicular to the nitrocellulose strip, with the shoot (scion) overhanging on the agar. A sharp cut was then performed in the first top mm of the hypocotyl. The hypocotyl stump (scion) and root (root stock) were grafted onto the relevant rootstock and scion, respectively, as indicated, by simple contact through the clean cuts, maintained by dint of water surface-tension and ensuring the cotyledon petioles were slightly above the agar surface. This generated controlled reciprocal heterografts (WT scion / transgenic rootstock, and conversely) and homografts (scion and root stock of the same genotype, but severed from different seedlings as in heterografts). The plates were resealed and returned to the growth chamber. Five days later (5 DAG, 5 days after grafting), the seedlings were transferred to large square plates (12 x 12 mm) filled with similar media but without sucrose. For the auxin experiment, 20 µM NPA (N-1-Naphthylphthalamidic acid in DMSO) or appropriate volume of DSMO solvent was supplied to the media, and the thickness of agar gel underneath the scion was removed so that the scion never touched it. All scion-rootstock combinations compared within an experiment were raised alongside each other within each replicated plate.

For the plants grown on soil, on 5 DAG, grafted seedlings were transferred to pots filled with seed raising mix supplemented with Osmocote slow release fertiliser (5g.L^-1^), and then raised alongside seedlings in plates, under the same standard conditions (12h daylength, 21°C, 120-140 µE measured at pot level).

### Microscopy

For *in vivo* confocal microscopy root, scion, or part of, were mounted on slides in ½ MS liquid media (0% sucrose, pH 5.7, 21°C) and imaged using a TCS-SP8 microscope (Leica, Germany) equipped with a 10X/0.3 NA or 63X/1.2 NA water immersion objective. Autofluorescence spectra were acquired on the same samples for spectral unmixing. Autofluorescence was excited using a 488 nm Argon laser and acquired in λ-mode (from 495 nm to 600 nm, 5 nm acquisition window) with the pinhole opened at 2.8 Airy unit. The same parameters were used to acquire GFP or YFP fluorescence on the same samples fluorescence and was spectra were subsequently unmixed using LAS-X (Leica) software. For monitoring the temporal and spatial patterns of appearance of scion-derived TCTP1-GFP fluorescence in the root, the whole primary root was imaged from 2 DAG, when graft junctions were strong enough, in x,y-mode. The fluorescence was acquired with a 10X/0.3 NA objective and a 510-525 nm acquisition window, with the same acquisition for all roots within an experiment, and each positive signal was confirmed by spectral unmixing.

For light microscopy, samples were mounted in water, and imaged in DIC (Differential Interferential Contrast) mode with a Leica DM5000B microscope fitted with a 40X/0.85 NA dry objective and a Leica DFC 310FX camera Leica Instruments). Individual cell lengths were measured along epidermal files from the quiescent centre to the root differentiation zone, using a custom macro in Fiji (code available upon request). Mature cell length was estimated by the means of 10 consecutive cells in the root differentiation zone.

### Phenotyping

For determination of the kinetics of root elongation, plates were scanned daily from seed germination using a flatbed scanner (Epson Perfection 2450 photo, Seiko Epson, Japan) at a resolution of 600 dpi, 8-bits per channel and saved in Jpeg. The raw images were automatically pre-processed (cropped and aligned with “Linear Stack alignment” plugin) with a custom “auto align” macro in Fiji, and root lengths (*l*) were measured using RootTrace as described in[76] (code available upon request). At least 6 plants per condition with a complete trace were available for each of the compared genotypes and conditions, and thus used for subsequent analysis. Traces of the primary root were manually curated and erroneous data were manually corrected using the “segmented line” tool in Fiji software. Relative root elongation rate (*RER*; mm mm^-1^h^-1^) was computed for each root as:

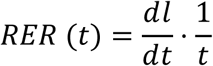

For normalization of average scion size between WT / RNAi and RNAi / WT grafts, 12 DAG i.e. a week after the grafted seedlings had been transferred onto a new plate as described above, the first leaf of WT scions was severed from the scion in half of WT / RNAi heterografts using a fine scalpel, generating WT_AMP_ scions, while the other half was left intact. To minimise potential artefacts due to confounding wounding effects when comparing amputated WT_AMP_ scions and scions left intact (WT or RNAi), one of the cotyledon stumps remaining from the cotyledons and severed from the latter on 0 DAG, at the time of grafting, was gently squeezed with a pair of very fine tweezers. All plates were scanned daily as previously described. Primary and secondary lateral root lengths were determined using Fiji software.

So that quantitative relationships between root system architecture and scion size could be examined, actual scion sizes were measured in all grafts at the end of each experiment through destructive sampling. Scions were severed from the rootstock and leaves were laid flat on a film of 0.8% phytagel, avoiding overlaps, and scanned with a flatbed scanner at a resolution of 2400 dpi, allowing for leaf area measurement using Fiji, and calculation of scion projected area. The roots were fixed in a fixing solution (Phosphate-buffered saline, 10% (v/v) Formaldehyde, 0.1% (v/v) Triton-X100), and imaged by light microscopy in DIC mode. Epidermal cell length profiles along the primary root were determined using the macro “Cell length profile”. The positions of all non-emerged lateral root primordia (*LR*P) and emerged lateral roots (*LR*) along the primary root were recorded through imaging the length of the whole root from hypocotyl junction to root tip. The distance between the most basal (oldest) lateral root and the most acropetal LRP was defined as the zone of lateral root formation and emergence. The densities of non-emerged and emerged roots along that zone could then readily be calculated.

### Molecular analysis

For qRT-PCR, fresh tissues were snap-frozen in liquid nitrogen and ground using a Tissue Lyzer (Quiagen). Total RNA was obtained using a chloroform/TRIzol (Thermofisher) extraction protocol, following manufacturer’s instructions. Messenger RNAs were purified from 40μg of total RNA using TYGR Dynabeads® oligo dT_25_ and subsequently used for cDNA synthesis using 200u of M-MLV (Promega), following manufacturer’s instructions. The qRT-PCR reaction was performed with 2.5uL of the diluted reverse transcriptase mix in a final volume of 10 μL of FastStart Universal SYBR® Green Master (Roche) and using a VIIA7 real time PCR system (Thermofisher). Primer efficiencies were calculated using LinRegPCR [77] and relative expression of target genes was normalised to four reference genes (*TIP41, APT1, UBC9 and PDF2*) using the delta-Ct method [78]. The primers used for quantification *of AtTCTP1* mRNAs are as described in [35]. GFP-specific primers were either GFP1_for_ 5’GATCCTGTTGACGAGGGTGT3’, GFP1_rev_ 5’GGATACGTGCAGGAGAGGAC3’, or GFP2_for_ 5’GATGCCGTTCTTTTGCTTGTCG and GFP2_rev_ 5’CGTGCAGTGCTTCTCCCGTTAC3’. The two sets of primers produced almost identical delta-Ct values, which were thus were averaged to calculate expression levels. Primers used for quantification of *CycB1;1* expression are CycB1;1_for_ 5’TCAGCTCATGGACTGTGCAA3’ and CycB1;1_rev_ 5’GATCAAAGCCACAGCGAAGC3’. In all experiments, replication consisted of 3 to 6 independent pools, each consisting of scion or rootstocks from at least 5 plants, unless specified otherwise.

### Statistical analysis

Data were analysed and plotted using the OriginPro software v9. This includes curve fittings, tests for equal variance (Levene) and ANOVA analysis of variance followed by post-hoc tests for pair-wise comparisons (Tukey or Bonferroni, as adequate).

### Gene accessions numbers

*AtAPT1* (*At1g27450*), *AtCyclinB-1* (*At4g37490*), *AtPDF2* (*At1g13320*), *AtTCTP1* (*At3g16640*), *AtTCTP2 (At3g05540), AtTIP41* (*At4g34270*), *AtUBC9* (*At4g27960*).

## Authors contribution

J.M. initiated the project; R.B. and J.M. conceived the experiments; R.B. performed the experiments; R.B. and J.M wrote the manuscript

## Data availability

The data supporting the findings of this study are available from the corresponding author upon reasonable request.

## Supplementary data

**Supplementary Figure 1.** TCTP1-GFP fluorescence is restricted to specific strands along the vasculature.

**Supplementary Fig. 2.** Distinctive GFP fluorescence pattern in roots grafted to a *pTCTP1∷gTCTP1-GFP* scion compared to a *p35S∷GFP* scion

**Supplementary Fig. 3**. Time course of appearance of TCTP1-GFP fluorescence in *pTCTP1∷gTCTP1-GFP* / WT heterografts.

**Supplementary Fig. 4.** Kinetics of scion expansion in homografts and heterografts between WT and TCTP-RNAi seedlings.

**Supplementary Fig. 5.** Similar scion and primary root sizes in WT_AMP_ and TCTP1-RNAi scions following scion size normalisation

**Supplementary Fig. 6.** Root patterning in TCTP1-RNAi seedlings shows no deviation from the stereotypical structure of WT Arabidopsis roots

**Supplementary Fig. 7.** At same scion size, primary and overall lateral root lengths are independent of TCTP1 expression in scion and mobility to rootstock

**Supplementary Fig. 8.** The auxin transport inhibitor NPA does not modify GFP fluorescence of scion-derived TCTP1-GFP protein in the root.

**Supplementary Table 1.** Detection of TCTP/TCTP-like messengers or proteins’ presence in the vasculature or movement through graft junctions, in published studies or databases.

