## Supplementary Figures for "Systemic signalling through *TCTP1* controls lateral root formation in Arabidopsis"

Supplementary Figure 1

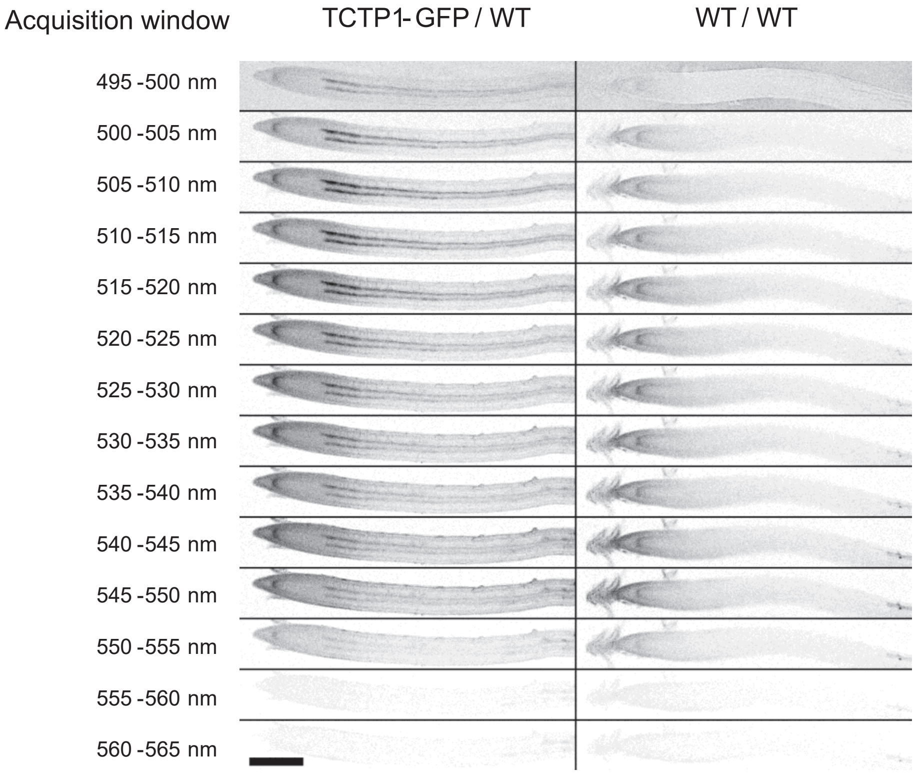


**Supplementary Figure 1. TCTP1-GFP fluorescence is restricted to specific strands along the vasculature.**

CLSM images of a WT primary root grafted onto a *pTCTP1::gTCTP1-GFP* scion (left panel), or WT scion (right panel). Images were acquired on 5nm emission windows between 495 nm and 560 nm, after excitation with a 488 nm laser. All frames were captured with the same microscope settings for the two roots. GFP-specific signal is restricted to two parallel strands, from root base to about 250 μm from the root tip. Maximum intensities between 505 and 520 nm correspond to the expected peaks for GFP emission. The two roots show autofluorescence at the quiescent centre. Scale bar, 200μm.

Supplementary Figure 2

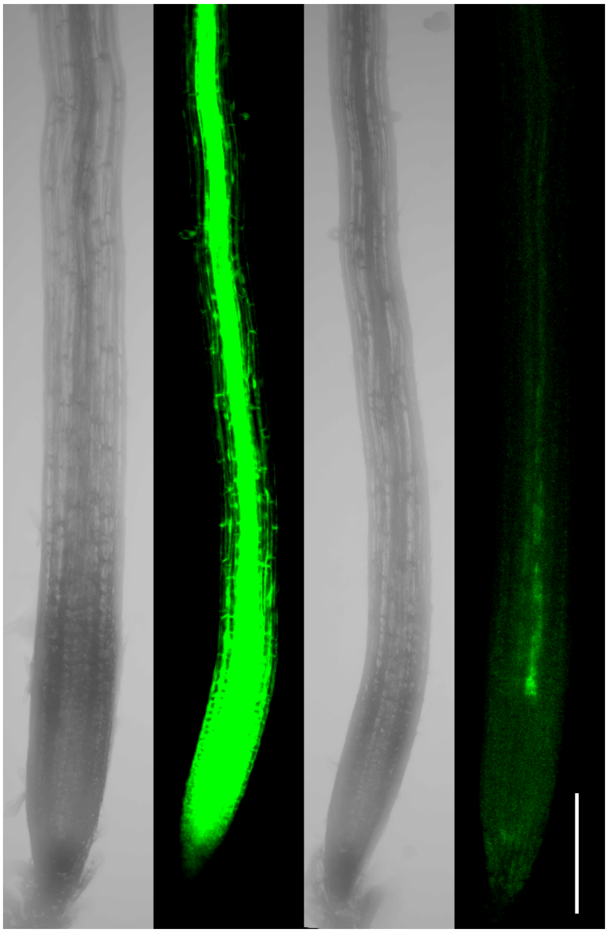


**Supplementary Fig. 2. Distinctive GFP fluorescence pattern in roots grafted to a *pTCTP1::gTCTP1-GFP* scion compared to a *p35S::GFP* scion**

Roots grafted to *35S::GFP* show diffuse fluorescence, with maximum intensity in the root meristem, consistent with passive diffusion as previously reported (Atkins et al. 2011; Paultre et al. 2016). Roots grafted to a *pTCTP1::gTCTP1-GFP* scion show faint fluorescence, restricted to thin strands along the vasculature, with no background autofluorescence in the meristem. Light transmitted channel images are also shown for the two roots. Scale bar, 200 µm

Supplementary Figure 3

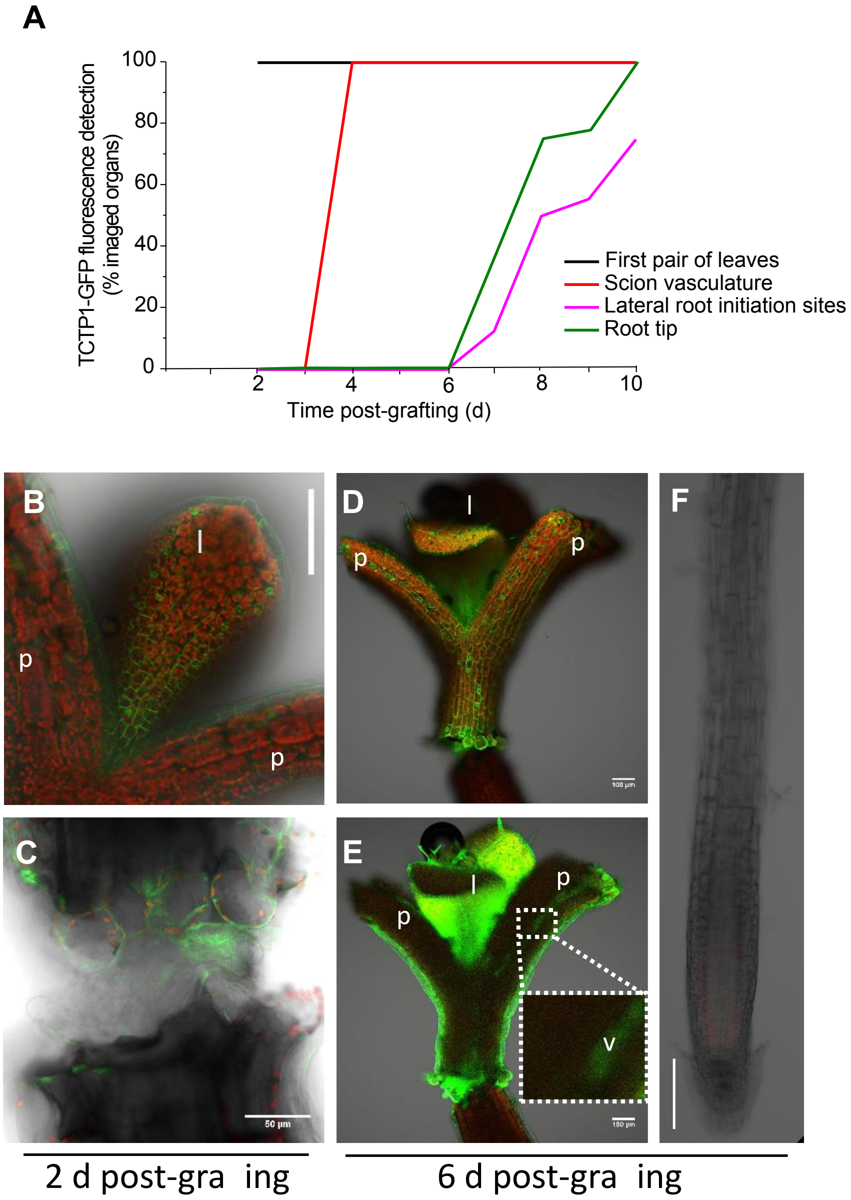


**Supplementary Fig. 3. Time course of appearance of TCTP1-GFP fluorescence in *pTCTP1::gTCTP1-GFP* / WT heterografts.**

**A**, Percentage of imaged seedlings where TCTP1-GFP fluorescence was detected in leaves, petiole vasculature, root tip and LR initiation sites through time (days after grafting, DAG), n ≥ 8. **B** to **F**, Confocal laser scanning microscopy images of TCTP1-GFP / WT heterografts captured 2 DAG (**B, C**), and 6 DAG (**D-F**), keeping the same microscope settings. **B-C**, weak GFP fluorescence in petioles and base of the first leaf (**B**); scion and root stock were attached but their vasculature were not connected yet (**C**). Scale bars, 100 µm in **B** and 50 µm in **C**. **D-E** Two images of the same scion at different z positions 6 DAG to highlight the presence of TCTP1-GFP fluorescence in the scion vasculature (inset in **E**). Scale bars, 100 µm. Note that the still fragile scion-root stock attachment broke upon mounting and cover slip pressure but was clean and neat before mounting. (**F**) Corresponding rootstock. TCTP1-GFP fluorescence is not yet detected in the primary root tip 6 DAG. *p*, petiole of cotyledons; *l*, leaf; *v*, vasculature. Scale bars, 100 µm.

Supplementary Figure 4

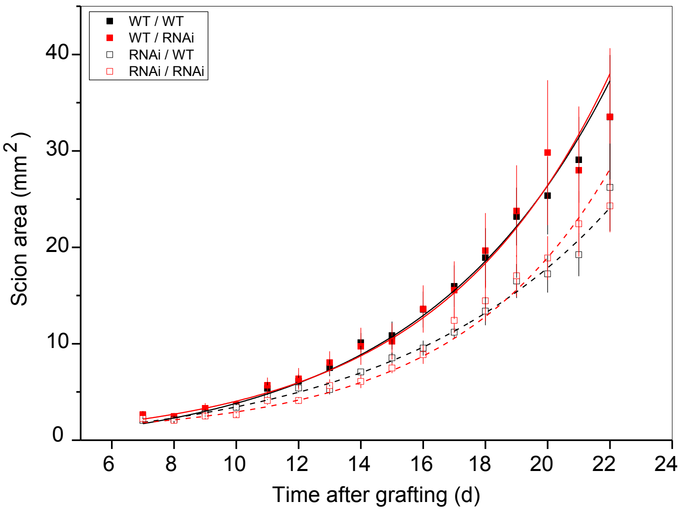


**Supplementary Fig. 4. Kinetics of scion expansion in homografts and heterografts between WT and TCTP-RNAi seedlings.**

Scion projected area as a function of time after grafting. Means ± SE, n = 5-8.
WT scion expansion rate was significantly higher than expansion rates of TCTP-RNAi scions, regardless of rootstock genotype.

Supplementary Figure 5

A

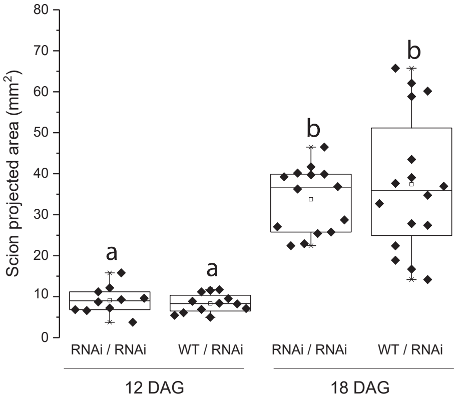


B

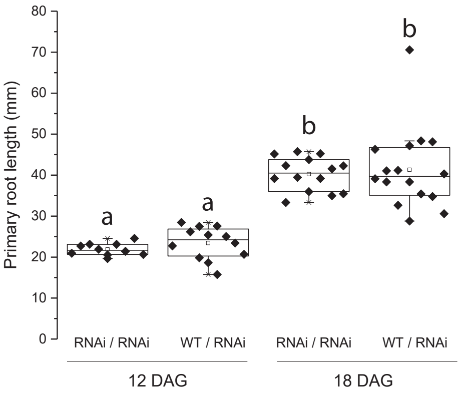


**Supplementary Fig. 5. Similar scion and primary root sizes in $WT_{AMP}$ and TCTP1-RNAi scions following scion size normalisation**

**A-B,** Scion projected area (**A**), and primary root length in RNAi/RNAi homografts and $WT_{AMP}$/RNAi heterografts 12 DAG and 18 DAG in an independent experiment from that shown with similar data in Fig. 4**G-I**. Box plots are as described in Fig. 4. Dots denote individual scions (**A**) and the associated roots (**B**); n = 10-16 grafted seedlings. Different letters denote statistically significant differences by two-way ANOVA followed by Bonferroni post-hoc test, *P* < 0.05. Scion sizes and primary root lengths were similar in the two sets of grafts, on both days.

Supplementary Figure 6

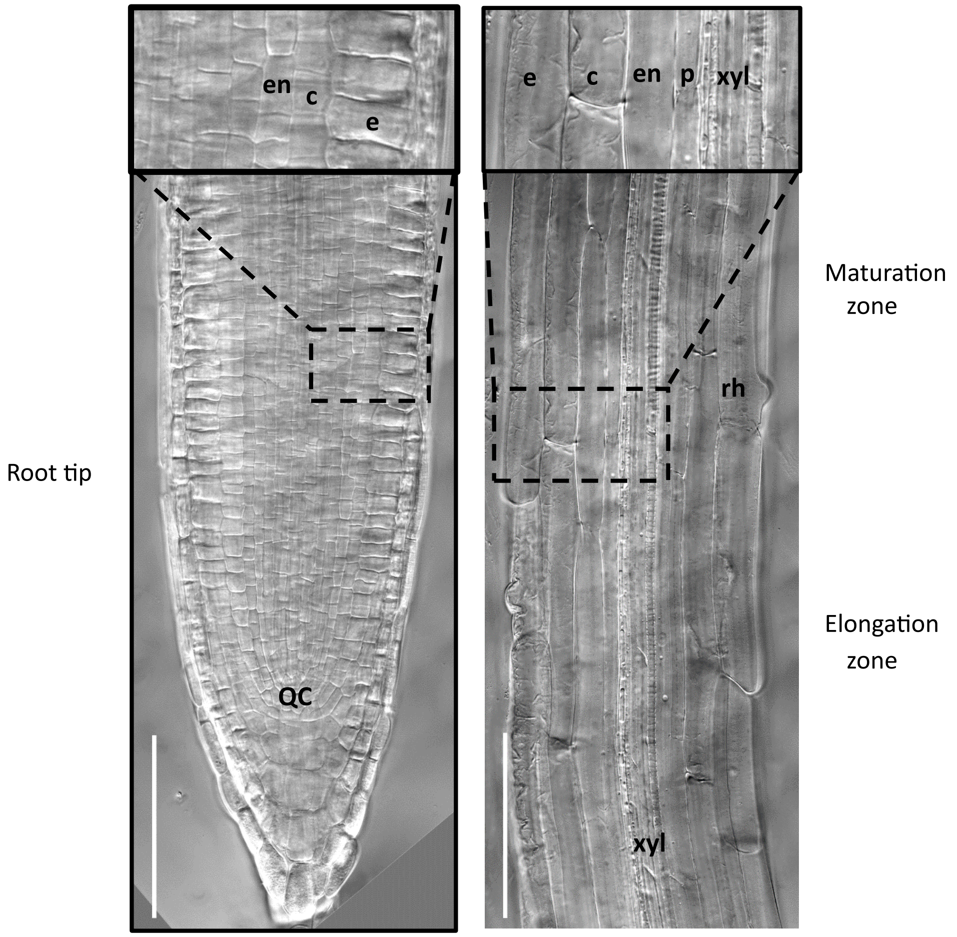


**Supplementary Fig. 6. Root patterning in TCTP1-RNAi seedlings shows no deviation from the stereotypical structure of WT Arabidopsis roots**

Frames captured by Differential Interference Contrast light microscopy. Root meristem and transition zone (left panel); Elongation and maturation (differentiation) zones (right panel). The insets show higher magnification enlargements for better visualisation of the root structure; epidermis (e), cortex (c), endodermis (en), pericycle (p), xylem (xyl). QC denotes the quiescent centre. Scales bars, 100 µm.

Supplementary Figure 7

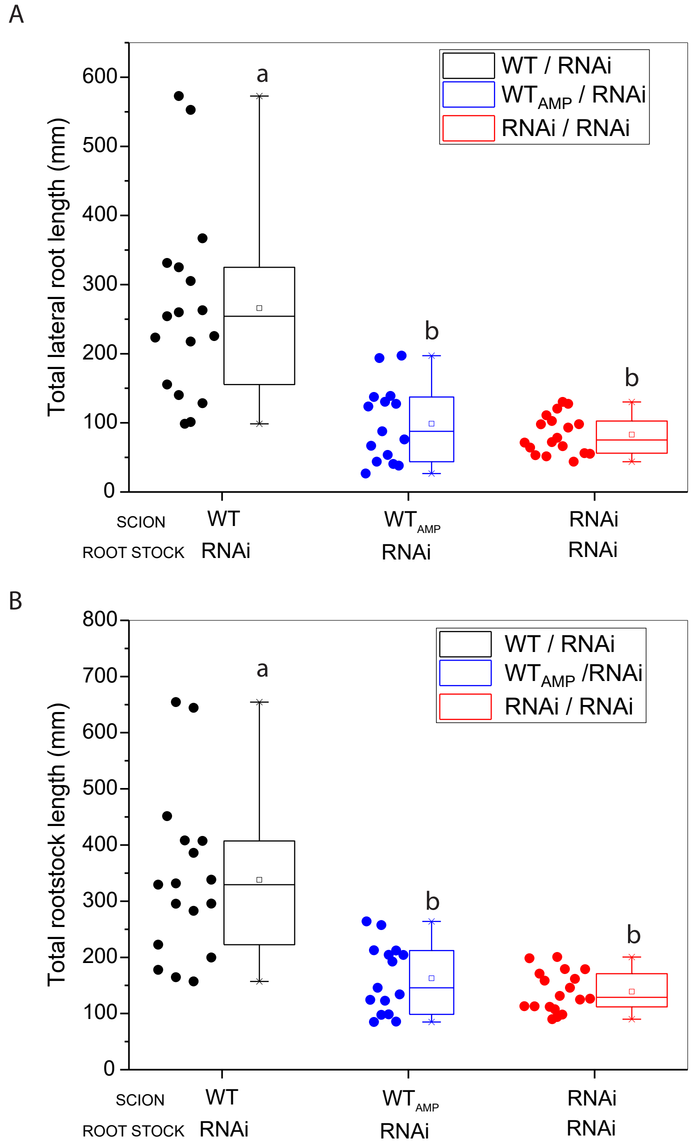


**Supplementary Fig. 7. At same scion size, primary and overall lateral root lengths are independent of *TCTP1* constitutive expression in scion and *TCTP1* mobile gene products to rootstock**

**A**, Cumulated lateral root lengths and total root length over the whole root system in WT / RNAi, WT$_{AMP}$ / RNAi, and RNAi / RNAi grafts, 22 DAG. Data points correspond to values for individual plants, n = 15-18 per scion/rootstock combination. Different letters denote statistically significant differences by one-way ANOVA followed by Bonferroni post-hoc test, $P < 0.05$.

### Supplementary Fig 8

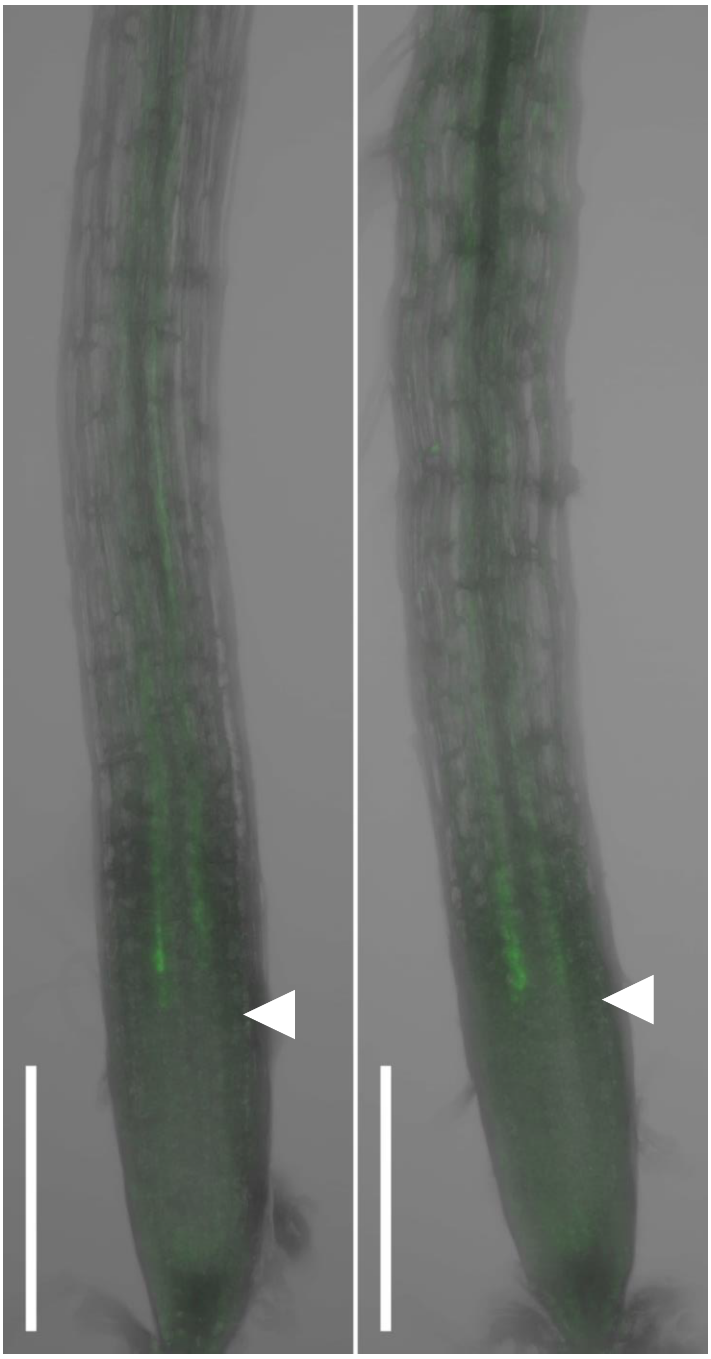


**Supplementary Fig. 8. The auxin transport inhibitor NPA does not modify GFP fluorescence of scion-derived TCTP1-GFP protein in the root.**

Maximal intensity projection of z-stack CLSM images representative of AtTCTP1-GFP fluorescence in primary root tips of 14 days old AtTCTP1-GFP / TCTP1-RNAi1 grafted plants, after 2 d exposure to DMSO (left) or 25 µM NPA (right). n ≥ 5 plants each. Green, AtTCTP-GFP fluorescence, grey, transmition. Scale bar, 200 um. The white arrow-head indicates the upper boundary of the root meristem.
