## Supplementary Table 1 for "Systemic signalling through *TCTP1* controls lateral root formation in Arabidopsis"

### Supplementary Table 1: Detection of TCTP/TCTP-like messengers or proteins presence in the vasculature, phloem or xylem sap or movement through graft junctions, in published studies or publicly available databases.

#### Mobility of TCTP/TCTP-like mRNAs through graft junctions

| Reference | Directionality of movement | *TCTP* gene ID or closest homolog | Plant species | Tissues analysed | Detection method |
|---|---|---|---|---|---|
| (Thieme et al., 2015) | host to parasite, Scion to rootstock | AT3G16640 | *A.thalianagrafts (Col-0 and Ped heterografts); A.thaliana(host) / C.reflexa (parasite)* | Grafted tissues and parasite | sequencing, RT-PCR |
| (Yang et al., 2015) | bidirectional | GSVIVG01017723001 | *vitis* spp | Grafted tissues | sequencing |
| (Kim et al., 2014) | host to parasite | AtTCTP1 | *A.thaliana(host) / C.reflexa (parasite)* | Parasite | sequencing |
| (Toscano-Morales et al., 2014) | bi directional but preferentially stock to scion | AtTCTP2 | *N.benthiamana* | Grafted tissues | RT-PCR and in situ hybridization |
| (Zhang et al., 2016) | Scion to sink (developing leaf), scion to root tip under normal and Phosphate inorganic deprivation | Csa3M154390 | *C.melo* / *C.lanatus* heterografts | Grafted tissues | sequencing |

### Presence of TCTP mRNA in phloem exudates

| Reference | TCTP reference or (closest) homolog | Plant species | Tissues used for sap collection | Sap collection method |
|---|---|---|---|---|
| (Deeken et al., 2008) | AT3G16640 [source] | A.thaliana | phloem exudate of Arabidopsis leaves | EDTA-exudation and LMPC-exudation |
| (Doering-Saad et al., 2006) | AM267384 [source] | R.communis | phloem exudate of hypocotyls | nd |
| (Deng et al., 2018) | HbTCTP/HbTCTP1 | H.brasiliensis | phloem exudates | nd |

**Curation of phloem sap proteome studies for presence/absence of TCTP/TCTP-like proteins.**

**Legend : C = cut (e.g. cut petiole, stem); I = incision(s); E = EDTA facilitated; S = stylectomy; DAE = Day after emergence; DAS = Day After Sowing ; w = weeks**

| Reference | Plant species | Method used for sap collection | Treatment | Age and / or tissue | TCTP presence | TCTP mentioned in the main text or supplementary data | TCTP reference or (closest) homolog | Comments on presence, abundance,... |
|---|---|---|---|---|---|---|---|---|
| (Aoki et al., 2005) | *C.maxima* | | Infected leaf | Leaves and roots | Yes | Main text | CmaCh11G012000 | |
| (Batailler et al., 2012) | *A.thaliana* | E | Healthy | Mature leaves (rosette, early and late flowering stages) | Yes | Main text | AtTCTP1 | |
| (Giavalisco et al., 2006) | *B.napus* | I | Healthy | 6-8 w inflorescence stems | Yes | Main text | Bol022977 (Relative) | Two isoforms reported with different molecular weight and isoelectric point |
| (Lin et al., 2009) | *C.maxima* | C | Healthy | 8 w old | Yes | Main text | CmaCh00G004010 | |
| (Fröhlich et al., 2012) | *C.maxima* | C | Healthy | 4-5 w stem/petioles | Yes | Supp data | not enough information | |
| (Malter and Wolf, 2011) | *C.melo* | C | Healthy/Infected (CMV infection) | 8-10 w old stems /petioles | No/Yes | Main text | MELO3C006670 | |
| (Barnes et al., 2004) | *R.communis* | I,S | Healthy | Mature leaf | Yes | Main text | not enough information | Two isoforms with same mass but different isoelectric point |
| (Aki et al., 2008) | *O.sativa* | S | Healthy | 5 w leaf | Yes | Main text | Os11g43900 aka Os11g0660500 | |
| (Rodriguez-Medina et al., 2011) | *L.albus* | I | Healthy | Flowering stage, collection from fruits, inflorescence, stem | Yes | Sup data | | |
| (Fan et al., 2015) | *C.sativus (NaCl sensitive)* | C | Healthy/NaCl stress | 2 w | Yes | Main text | not enough information | 80% reduction of TCTP abundance in phloem sap after NaCl stress |
| | *C.sativus (NaCl tolerant)* | | | | Yes | Main text | not enough information | |
| (Du et al., 2015) | *O.sativa cv 9311 (Susceptible)* | E | Healthy/Insect (Brown plant Hopper) | 4-leaf stage | Yes | Main text | Os11g43900 aka Os11g0660500 | |
| | *O.sativa cv B5 (Resistant)* | | | | Yes | Main text | | |
| (Carella et al., 2016) | *A.thaliana* | E | Healthy | 4 w leaf | Yes | Supp data | AtTCTP1 | Increase or decrease of AtTCTP1 abundance after infection with avirulent or virulent P.syringae, respectively |
| | | | Infected with virulent P.syringae(PSTDC3000) | | Yes | | | |
| | | | Infected with avirulent P.syringae (avRpt2) | | Yes | | | |
| (Walz et al., 2002) | *C.maxima* | C | Healthy/Drought | 6-8 w adult stems | ND / ND | | | |
| | *C.melo* | | | | ND / ND | | | |

| | | | | | |
|---|---|---|---|---|---|
| (Cho et al., 2010) | *C.maxima* | C | Healthy | 2 months stems | ND |
| (Gaupels et al., 2008) | *H.vulgaris* | S | Healthy | 8-12 d after germination | ND |
| (Guelette et al., 2012) | *A.thaliana* | E | Healthy | 6 w rosette leaves | ND |
| (Lattanzio et al., 2013) | *L. texensis* | I | Healthy | Inflorescence base | ND |
| (Dafoe et al., 2009) | *P.trichocarpa* | E | Healthy/Wounded | 3 months 5LPI index = 5-16 | ND / ND |
| (Gaupels et al., 2012) | *C.maxima* | C/S | Healthy / Wounding | 4-5 w petioles and stems | ND / ND |
| (Gutierrez-Carbonell et al., 2015) | *B.napus* | I | Healthy / Fe-deficiency | 8 w inflorescence | ND / ND |
| (Serra-Soriano et al., 2015) | *C.melo* | C | Healthy / infected with virus (MNSV-Al) | 6-10 d | ND / ND |

**Curation of xylem sap proteome studies for presence/absence of TCTP/TCTP-like proteins.**

**Legend : C = cut (e.g. cut petiole, stem); F = forced pressure; DAE = Day after emergence ; DAS = Day After Sowing ; w = weeks**

| Reference | Plant species | Method used for sap collection | Treatment | Age | TCTP presence | TCTP mentioned in the main text or supplementary data | TCTP reference or (closest) homolog | Comments on presence, abundance |
|---|---|---|---|---|---|---|---|---|
| (Gawehns et al., 2015) | *S.lycopersicum* | C | Healthy/Infected with Fusarium | 6 w | Yes | Main text | Solyc01g099770 | Increased relative abundance after infection |
| (Abeysekara and Bhattacharyya, 2014) | *G.max* | CF | Healthy/Infected with *F.virguliforme* | 2-3 w | Yes | Supplementary data | GLYMA09G04950 | detected more often in infected tissues |
| (Buhtz et al., 2004) | *C.sativus* | C | Healthy | 8w | ND | | | |
| (Buhtz et al., 2004) | *C.maxima* | C | Healthy | 8 w | ND | | | |
| (Buhtz et al., 2004) | *B.napus* | C | Healthy | 8-10 w | ND | | | |
| (Buhtz et al., 2004) | *B.oleracea* | C | Healthy | 8-10 w | ND | | | |
| (Kehr et al., 2005) | *B.napus* | C | Healthy | 12 w, flowering | ND | | | |
| (Ligat et al., 2011) | *B.oleracea* | C | Healthy | 6-8 w old | ND | | | |
| (Djordjevic et al., 2007) | *G.max* | CF | Healthy | 1 month | ND | | | |
| (Krishnan et al., 2011) | *G.max* | CF | Healthy | ND | ND | | | |
| (Zhang et al., 2015) | *G.hirsutum* | C | Healthy | 100 DAE, Flowering stage | ND | | | |
| (Aki et al., 2008) | *O.sativa* | C | Healthy | 5 w | ND | | | Same authors reported TCTP in phloem |
| (Agüero et al., 2008) | *V.vinifera* | C | Healthy | 2 years old and adult | ND | | | |
| (Alvarez et al., 2006) | *Z.mais* | CF | Healthy | 23 DAS | ND | | | |
| (Rep et al., 2002) | *S.lycopersicum* | C | Healthy/Infected (*F.oxysporum* ) | 5 – 8 w old | ND/ND | | | |

| | | | | | |
|---|---|---|---|---|---|
| (Houterman et al., 2007) | *S.lycopersicum* | C | Healthy/Infected (*F.oxysporum* ) | 7 w old | ND/ND |
| (Pu et al., 2016) | *B.oleracea* | C | Healthy/Infected (*F.oxysporum* ) | 6 w old | ND/ND |
| (Subramanian et al., 2009) | *G.max* | C | Healthy/infected (*B.japonicum* ) | 1 w old | ND/ND |
| (Floerl et al., 2008) | *B.napus* | CF | Healthy/Infected (*V.Longisporum* ) | more than 2 w | ND/ND |
| (Liao et al., 2012) | *Z.mais* | CF | Healthy/ Nitrogen supply | V12 stage | ND/ND |
| (Fernandez-Garcia et al., 2011) | *B.oleracea* | CF | Healthy/ NaCl stress | 33 d | ND/ND |
